# Single-molecule analysis of specificity and multivalency in binding of short linear substrate motifs to the APC/C

**DOI:** 10.1101/2021.09.25.461797

**Authors:** Nairi Hartooni, Jongmin Sung, Ankur Jain, David O. Morgan

## Abstract

Robust regulatory signals in the cell often depend on interactions between short linear motifs (SLiMs) and globular proteins. Many of these interactions are poorly characterized because the binding proteins cannot be produced in the amounts needed for traditional methods. To address this problem, we developed a single-molecule off-rate (SMOR) assay based on microscopy of fluorescent ligand binding to immobilized protein partners. We used it to characterize substrate binding to the Anaphase-Promoting Complex/Cyclosome (APC/C), a ubiquitin ligase that triggers chromosome segregation. We find that SLiMs in APC/C substrates (the D box and KEN box) display distinct affinities and specificities for the substrate-binding subunits of the APC/C, and we show that multiple SLiMs in a substrate generate a high-affinity multivalent interaction. The remarkably adaptable substrate-binding mechanisms of the APC/C have the potential to govern the order of substrate destruction in mitosis.

## Introduction

Inside the crowded and noisy confines of the cell, clear and robust regulatory signals require highly specific protein-protein interactions. Many of these interactions depend on the binding of a globular domain in one protein to short linear sequence motifs (SLiMs) in another. SLiMs are short conserved amino acid sequences that are generally found in disordered protein regions, and a remarkably diverse variety of SLiMs are involved in numerous regulatory processes^1^. The affinities and specificities of SLiMs for their targets determine the impact of these motifs in signaling, but we have only a limited understanding of these interactions.

The central importance of SLiM interactions is illustrated by substrate binding to the Anaphase-Promoting Complex/Cyclosome (APC/C)^2^. The APC/C is a conserved 13-subunit ubiquitin ligase that triggers the destruction of key proteins controlling the initiation of chromosome segregation in mitosis^3–6^. Its substrates include the separase inhibitor securin, whose destruction allows separase to separate the duplicated chromosomes. Another key APC/C target is mitotic cyclin, whose destruction is required for late mitotic events. Disordered regions in these substrates contain SLiMs, or degrons, that bind to specific subunits of the APC/C. The APC/C holds the substrate in place while an E2 co-enzyme binds nearby and transfers ubiquitin to a lysine on the substrate or on ubiquitin. Repeated ubiquitin transfer from multiple E2s leads to the formation of polyubiquitin chains that are recognized by the 26S proteasome, resulting in substrate degradation.

The APC/C is activated in mitosis by one of two related substrate-binding subunits called Cdc20 and Cdh1. These activators contain a globular WD40 domain that binds substrate degrons, flanked by partially disordered regions that mediate binding to the APC/C, resulting in a conformational change that enhances binding of the E2 co-enzyme^7, 8^. Activators interact transiently with the APC/C at specific cell cycle stages. Cdc20 activates the APC/C during metaphase and anaphase of mitosis and binds a narrow range of substrates governing the initiation of chromosome segregation^9^. In late anaphase, Cdc20 is replaced by Cdh1, which activates the APC/C in late mitosis and G1^9^. Cdh1 has broader specificity and targets many additional proteins for destruction.

Three major degrons have been identified in APC/C substrates: the destruction box (D box), KEN box, and ABBA motif^2,10–16^ (Fig. 1a). As with most SLiMs, these degrons are found in disordered regions, and substrates often contain multiple degrons. The most important degron is the D box, which has a composite binding site involving both the WD40 domain of the activator and the Apc10/Doc1 subunit of the APC/C. The conserved residues of the D box are RxxLxxxxN. The N-terminal RxxL segment interacts with an acidic patch and aliphatic pocket on the WD40 domain of the activator^10^. The C-terminal residues of the D box interact with the Apc10 subunit^16–18^. As a result, the D box helps anchor the activator to the APC/C^19, 20^. The second major APC/C degron, often found near a D box, is the KEN box, which usually contains a well-conserved KEN sequence that interacts with a specific binding pocket on the activator WD40 domain^2, 10^. Lastly, the less common ABBA motif has a complex consensus sequence that interacts with a specific groove on the activator WD40 domain^2, 11^. In yeast, variations in this motif result in specificity for one or the other activator^2, 11, 15^.

**Fig. 1:**
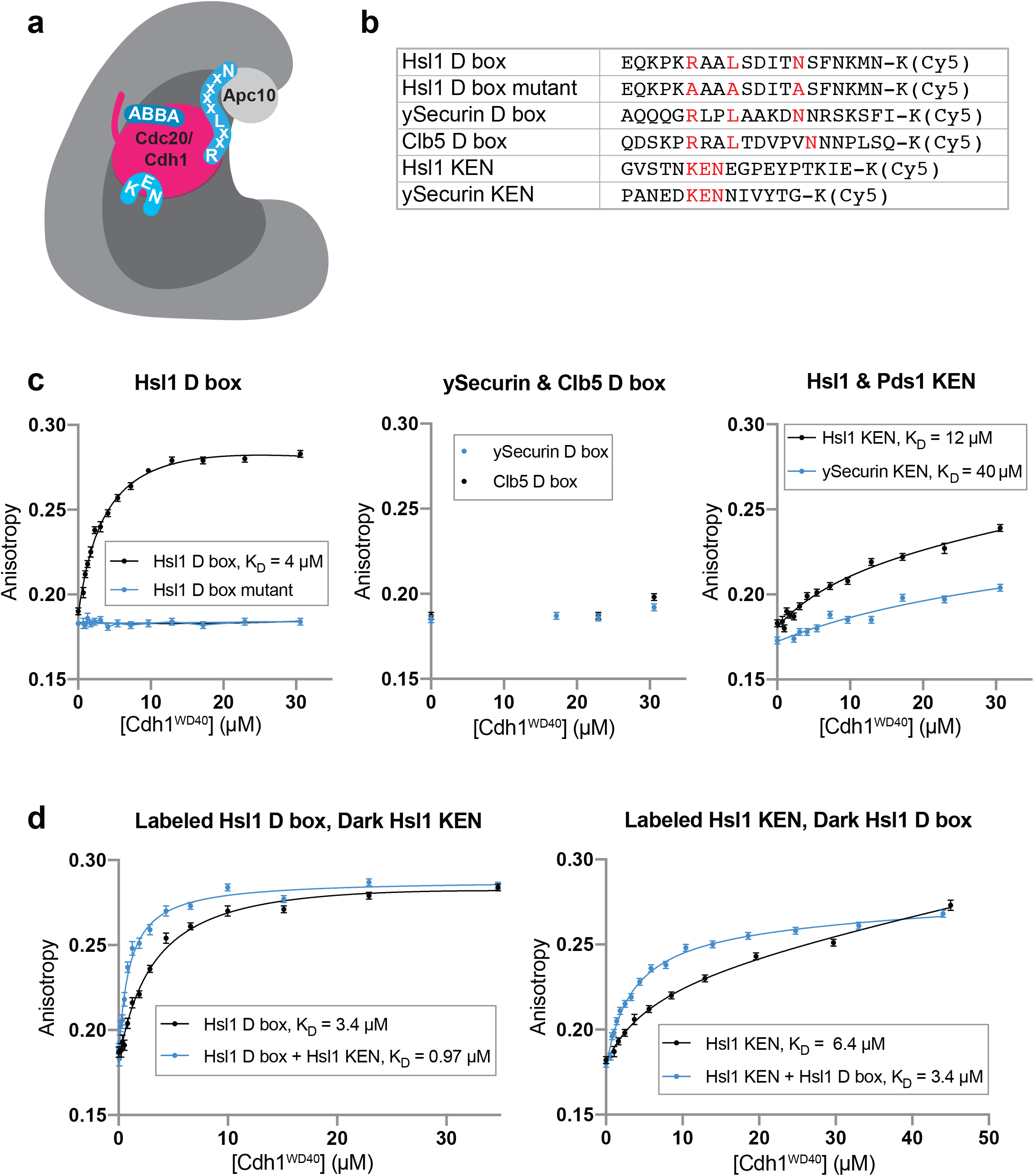
Analysis of degron affinity by fluorescence anisotropy. **a**, Cartoon summarizing the interactions of activated APC/C with the three major degrons found in APC/C substrates. **b**, List of Cy5-labeled degron peptides tested in anisotropy experiments. **c**, Results of fluorescence anisotropy experiments performed using the peptides listed in **b**. 10 nM peptide was incubated with up to 30.6 µM Cdh1^WD40^. Data points represent mean +/- SD (n = 10 reads per reaction). Data is representative of two independent experiments. **d**, Binding was measured with labeled degron peptide (10 nM) in the absence (black lines) or presence (blue lines) of an unlabeled version of the other degron (100 µM). Data points represent mean +/- SD (n = 10 reads per reaction).

APC/C substrates are targeted for destruction in a specific order during mitosis. Substrates that control anaphase onset, such as securin, are generally degraded earlier, in metaphase, than substrates involved in late mitotic events. This order is likely to be achieved in part by selectivity of the activators for different substrates. Destruction of securin and a small number of other early substrates depends on Cdc20, whereas numerous later substrates, degraded in late mitosis and G1, are targeted specifically by Cdh1^19, 21^. There is some evidence for activator-specific D-box sequences, as well as evidence that the KEN box has a preference for Cdh1^12^. However, activator specificity alone cannot explain all substrate ordering. The same activator is known to target different substrates at different times, perhaps due to variations in degron affinity, combinations of multiple degrons, or other mechanisms^2, 15, 22, 23^.

Substrate affinity for the APC/C is a critical determinant of the extent of ubiquitylation. As a ubiquitin ligase of the RING family, the APC/C binds substrates at one site while the E2-ubiquitin conjugate binds at a nearby site, enabling lysines in the disordered substrate to attack the E2 to catalyze transfer^3, 4, 6^. Ubiquitylation is processive: multiple E2-ubiquitin conjugates can bind, transfer ubiquitin, and dissociate during a single substrate-binding event^24–27^. Thus, the number of ubiquitins added is directly dependent on substrate dwell time, or dissociation rate. It is likely that proteasome recognition depends on the number and length of polyubiquitin chains, so different substrate dwell times are likely to influence the timing of their degradation^22, 28^.

Despite decades of research on the APC/C, the affinity of substrate binding remains poorly understood. Conventional approaches to affinity analysis are hampered by our inability to express and purify large amounts of the multi-subunit APC/C or its activators. To solve this problem, we developed a single-molecule binding assay that provides robust measurements of the rate of dissociation of substrates bound to activators and the activated APC/C. These methods provide important new insights into degron affinity, activator specificity, and multivalency. Our methods can also be applied to binding interactions with other proteins and protein complexes that are not readily studied by conventional ensemble methods.

## Results

### Analysis of degron affinity by ensemble biochemistry

To determine the affinities of APC/C degrons for their binding sites, we first used conventional equilibrium binding assays of degron peptide binding to the activator. We used the baculovirus system to produce the WD40 domain of Cdh1 from the budding yeast *Saccharomyces cerevisiae* and measured fluorescence anisotropy to assess the binding of fluorescently-tagged degron peptides at increasing activator concentrations. The WD40 domain of yeast Cdc20 could not be expressed and was not studied.

Our studies centered on the well-known D box of the yeast protein Hsl1 (Fig. 1b), a late mitotic substrate that is targeted primarily by APC/C^Cdh1^ in vivo^29^. The Hsl1 D box is known to have an ideal consensus sequence that binds tightly to APC/C^Cdh1^, resulting in highly processive modification^19, 30^. We found that the Hsl1 D box binds the Cdh1 WD40 domain with a dissociation constant (K_D_) of 4.0 µM (Fig. 1c). Binding was abolished by mutation of three key residues in the D box.

We also analyzed the D boxes of yeast securin/Pds1(ySecurin) and the S-phase cyclin Clb5. These proteins are targeted by APC/C^Cdc20^ prior to anaphase in vivo but are also thought to be modified by APC/C^Cdh1^ in G1^15^. Neither D box displayed significant binding to Cdh1 in our assay, suggesting that these D boxes have low affinity for this activator (Fig. 1c).

We also tested the KEN boxes of Hsl1 and ySecurin. Both peptides bound with low affinity, such that binding saturation was not achieved at 30.6 µM Cdh1, but we obtained reasonable estimates of 12 µM and 40 µM for the K_D_ values of the Hsl1 and ySecurin KEN degrons, respectively (Fig. 1c). We also tested a Cdh1-specific ABBA motif from the pseudosubstrate yeast protein Acm1. This motif bound with very low affinity to Cdh1 (Supplementary Data Fig. 1).

Substrates of the APC/C often contain both a D box and a KEN box, generally at a distance that should allow simultaneous binding. We tested the possibility that binding of one degron affects affinity for the other. Addition of saturating unlabeled Hsl1 KEN peptide improved affinity for the Hsl1 D box about 3-fold (Fig. 1d). As expected for an allosteric mechanism, the reverse was also true: saturating D box peptide improved affinity for the KEN box about 2-fold (Fig. 1d).

### APC/C single molecule assay development

Conventional assays like that used in Fig. 1 are limited by the need for very large amounts of purified binding protein, which is possible for the Cdh1 WD40 domain but not possible for Cdc20 or for the APC/C or APC/C-activator complexes. To thoroughly probe the interaction of substrates with the APC/C, we therefore developed a Single Molecule Off Rate (SMOR) assay in which dynamic substrate-APC/C interactions can be visualized and quantified by fluorescence microscopy. Our goal was to create an adaptable and simple-to-use platform that could be deployed to probe protein-protein interactions for any protein or protein complex that cannot be purified in large quantities.

We applied a previously developed antibody-based method to tether single protein molecules on a functionalized glass surface^31^, and modified it to capture transient interactions with fluorescent ligands using total internal reflection fluorescence (TIRF) microscopy.

NeutrAvidin and biotinylated antibody were used to tether molecules to glass cover slips in small chambers with ports for influx and outflow of ligand solutions^31^. We populated the surface with budding yeast activator (Cdh1^WD40^), activated APC/C (APC/C^Cdh1^ or APC/C^Cdc20^), or APC/C lacking activator (APC/C^apo^). Activator proteins were produced with the baculovirus system, APC/C was purified from yeast cells, and APC/C-activator complexes were prepared by mixing purified APC/C and activator prior to immobilization on the glass (Supplementary Data Fig. 2a). Very small amounts of protein were required. Typically, excellent glass coverage could be achieved with less than a nanogram of protein.

To confirm successful capture of the target molecule on the glass surface, C-terminal GFP tags were fused to the Cdh1^WD40^ protein and the Apc1 subunit of the APC/C. The C-terminal Apc1 tag did not affect APC/C ubiquitylation activity in vitro (Supplementary Data Fig. 2b). Activators were N-terminally tagged with a Strep Tag II, which had no effect on ubiquitylation activity in vitro (Supplementary Data Fig. 2c). Proteins were immobilized on glass using either biotinylated anti-Strep Tag II antibody to bind activator or anti-GFP antibody to bind the GFP-tagged APC/C^apo^ (Fig. 2a).

**Fig. 2:**
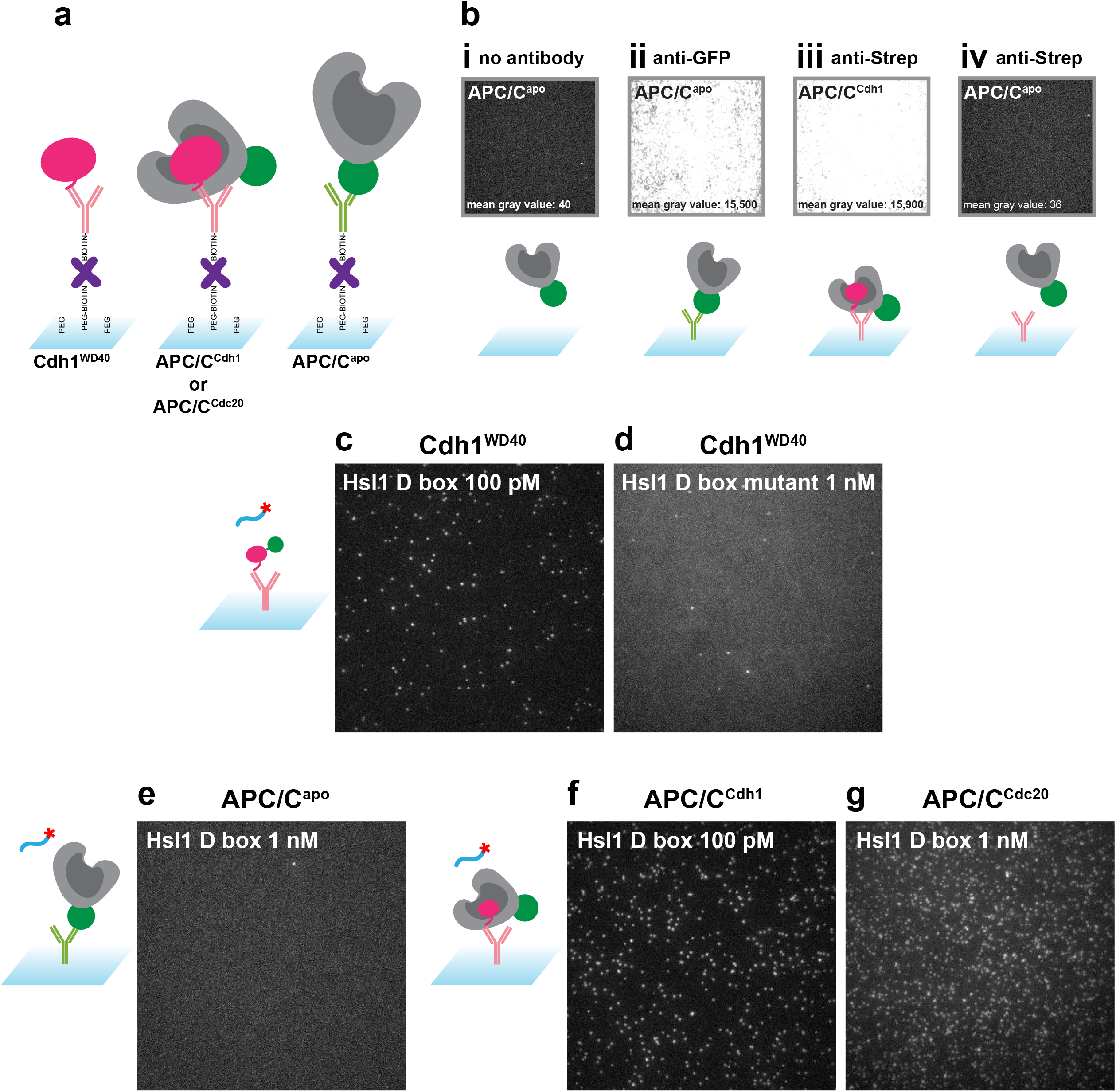
SMOR assay setup. **a**, Immobilization of binding proteins on cover glass surface. For activator or for activated APC/C, we used biotinylated anti-Strep-Tag II antibody (pink), which binds the Strep-Tag II on the activator N-terminus. For APC/C^apo^, we used biotinylated anti-GFP antibody (green), which binds the GFP tag on the Apc1 subunit. Antibody is linked to the biotinylated PEG on the surface using NeutrAvidin (purple). **b**, Fluorescent signal from APC/C-GFP in the absence and presence of antibodies and activator as indicated. **c**, **d**, Single-molecule interactions of Cy5-labeled Hsl1 D box peptide (**c**) or mutant peptide (**d**) with immobilized GFP-tagged Cdh1^WD40^. **e**, Lack of interactions between Hsl1 D box peptide and immobilized APC/C^apo^. **f**, **g**, Single-molecule interactions of Cy5-labeled Hsl1 D box peptide with immobilized APC/C^Cdh1^ (**f**) or APC/C^Cdc20^ (**g**). Images in **c**-**g** are maximum intensity projections of the first 10 frames of a movie at continuous exposure and 100 ms frame rate.

GFP fluorescence was not observed when there was no antibody immobilized on the surface, indicating that our glass functionalization scheme minimized background APC/C binding (Fig. 2bi). In contrast, anti-GFP antibody specifically immobilized abundant APC/C^apo^ (Fig. 2bii). Similarly, anti-Strep Tag II antibody specifically immobilized activated APC/C, and no cross-reactivity with APC/C^apo^ was observed (Fig. 2biii and iv). Note that immobilization of activated APC/C with a tag on the activator subunit ensures that we are measuring interactions only with intact APC/C that retains activator binding activity.

We carried out initial binding studies with the same Cy5-labeled Hsl1 D box peptide that we used for our binding analysis in Fig. 1. Capturing the signal from the Cy5 dye by TIRF microscopy, we observed binding to Cdh1^WD40^ at 100 pM peptide (Fig. 2c), and very little binding with just antibody on the surface (Supplementary Data Fig. 3). The mutant D-box peptide displayed negligible binding (Fig. 2d). Furthermore, the Hsl1 D box peptide did not interact with APC/C^Apo^ or the anti-GFP antibody used to tether it to the surface (Fig. 2e, Supplementary Data Fig. 3). Although the Apc10 subunit of the APC/C is believed to interact with the C-terminal residues of the D box, the affinity of this interaction is known to be extremely low. Finally, we demonstrated that the Hsl1 D box peptide interacts with APC/C activated with either Cdh1 or Cdc20 (Fig. 2f, g).

### Computational analysis of ligand dwell time

To quantify the affinity of substrate-APC/C interactions, we next developed data analysis methods to determine the length of time that a fluorescent ligand remains bound to its binding partner on the glass surface. The reciprocal of the mean dwell time is a reasonable estimate of the dissociation rate constant, k_off_, which provides important clues about the extent of multiubiquitylation by the APC/C, and thus the substrate degradation rate in the cell^32^.

Signal intensity in single-molecule studies depends on the nature of the dye, the parameters of the microscope, and whether the light being captured is from a monomeric molecule or a much brighter multimer. Our analysis pipeline is designed to account for all these factors for both short and long binding events and is robust to experimental and technical perturbations. In short, the pipeline corrects movies for different intensities across the field of view, corrects for drift if needed^33^, and then analyzes information on signal intensity to identify single molecule binding events and calculate dwell time (Fig. 3a).

**Fig. 3:**
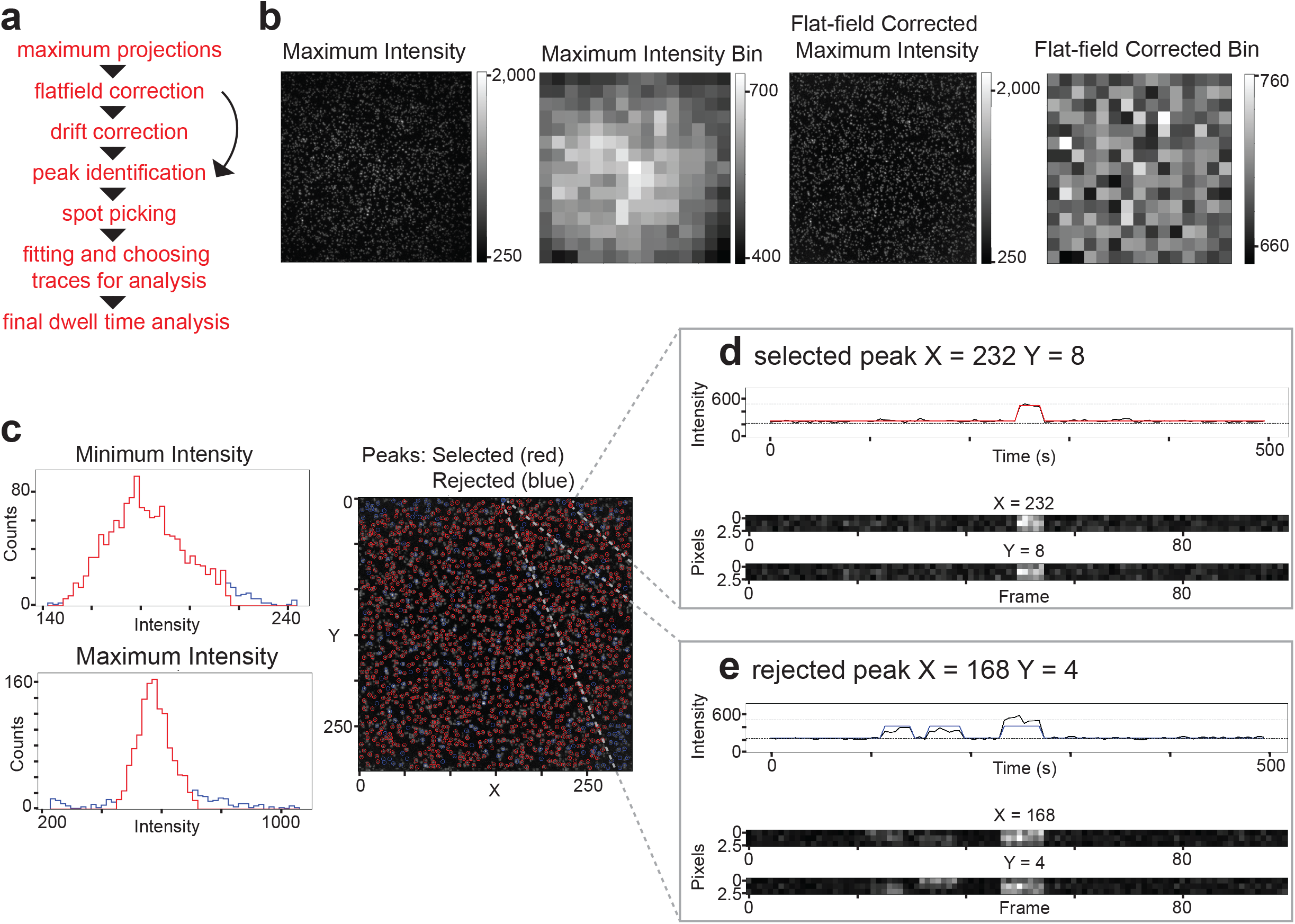
SMOR analysis. **a**, Analysis pipeline for SMOR analysis. **b**, For flatfield correction, the maximum intensity projection of a 300 x 300 pixel movie is binned into a 20 x 20 pixel grid, flatfield corrected, and then binned again to check correction. **c**, Histograms of minimum and maximum intensity values along traces for each peak. Red indicates selected peaks (≤3 SD from median) and blue indicates rejected peaks. At right is a maximum intensity projection with selected (red circle) and rejected (blue circle) peaks (x,y coordinates). **d**, Representative single-molecule trace of a selected peak at coordinates (232, 8). **e**, Representative single molecule trace of a rejected peak at coordinates (168, 4).

Fig. 3 illustrates our analysis methods using the binding of Cy5-labeled Hsl1 D box peptide to Cdh1-activated APC/C. First, the 512 x 512 pixel movie was cropped to the central 300 x 300 pixel grid. Analysis of signal intensity across the grid was then used to generate image projections of maximum and minimum intensities at all pixels during the length of the video (Fig. 3b and Supplementary Data Fig. 4a). The minimum intensity projection reveals a low level of long-lived nonspecifically bound fluorescent substrate on the glass. The maximum intensity projection shows potential transient binding events. In this example, there was a 10-fold difference between the range of intensities found in the maximum and minimum intensity projections (Supplementary Data Fig. 4a). Greater differences in intensity between the two indicates higher signal-to-noise ratio and thus more robust detection of binding events.

Binding signals in the maximum intensity projection tend to be brighter at the center of the TIRF evanescent wave, which can be a problem as we use the intensity level of the fluorescence signal to identify single molecules and discard multimers. To apply flat-field correction, intensities from the maximum intensity projection were used to create a mask of areas with any signal (Supplementary Data Fig. 4b). After application of the mask, the average intensities in 20 x 20 pixel grids were used to create an intensity bin image that shows the center of the TIRF evanescent wave (Fig. 3b). The edges of the bins were smoothened with a Gaussian filter to obtain an intensity bin filter, which was used to normalize intensities across the grid for flat-field correction (Supplementary Data Fig. 4b). To confirm flat-field correction, the same process was repeated on the corrected image, revealing more evenly distributed illumination (see Supplementary Methods). The corrected dataset was then applied to the next step, which for longer acquisition intervals includes drift correction (Supplementary Data Fig. 4c). Peak intensities or “peaks” were identified as 3 x 3 pixel squares (1 pixel = 16 x 16 µm) centered on (x, y) coordinates on this corrected maximum intensity projection (Supplementary Data Fig. 4d).

Next, we identified peaks that were most likely to represent genuine binding events. The first step was to discard peaks that were too bright, indicating a multimer. We created histograms of minimum and maximum intensities at each peak on the 300 x 300 pixel grid (Fig. 3c). Multimers were excluded in most cases by discarding peaks that were three standard deviations from the median maximum intensity. The same was done for minimum intensity peaks to eliminate background noise from dimmer signals.

For each selected peak, the signal intensity trace over time was fit to a Hidden Markov Model (HMM)^34^ to determine a bound/unbound state trajectory. First, the intensities throughout the movie along single molecule traces at all peaks were plotted in a histogram and fit using a double Gaussian to determine the mean maximum and minimum intensity values for the overall signal unique to each movie (Supplementary Data Fig. 4e). These initial parameters were used in the HMM to define bound and unbound states (Supplementary Data Fig. 4f, g). Deviation of the intensity data from the HMM fit for each trace was calculated using root mean squared deviation (RMSD). A histogram of RMSD values for HMM fitting of all traces was created, and a trace was rejected if the RMSD value was more than two standard deviations away from the median (Supplementary Data Fig. 4h). If a trace fell within the intensity parameters and had a low RMSD value, it was included in the analysis and colored red (Fig. 3d). If a trace had a high RMSD value, it was not included and colored blue. In the example in Fig. 3e, fluorescence at a nearby binding event created deviations in intensity during the movie and resulted in a higher RMSD value.

For each peak selected as a genuine binding event in the HMM, we used maximum likelihood estimation with an exponential distribution to statistically infer the dwell time. For each movie, a histogram showing the distribution of dwell times from multiple traces was calculated from the estimated inverse cumulative density function (Fig. 4). Note that oxygen scavenging agents were used in all experiments, ensuring that photobleaching occurred over much longer time scales than observed dwell times.

**Fig. 4:**
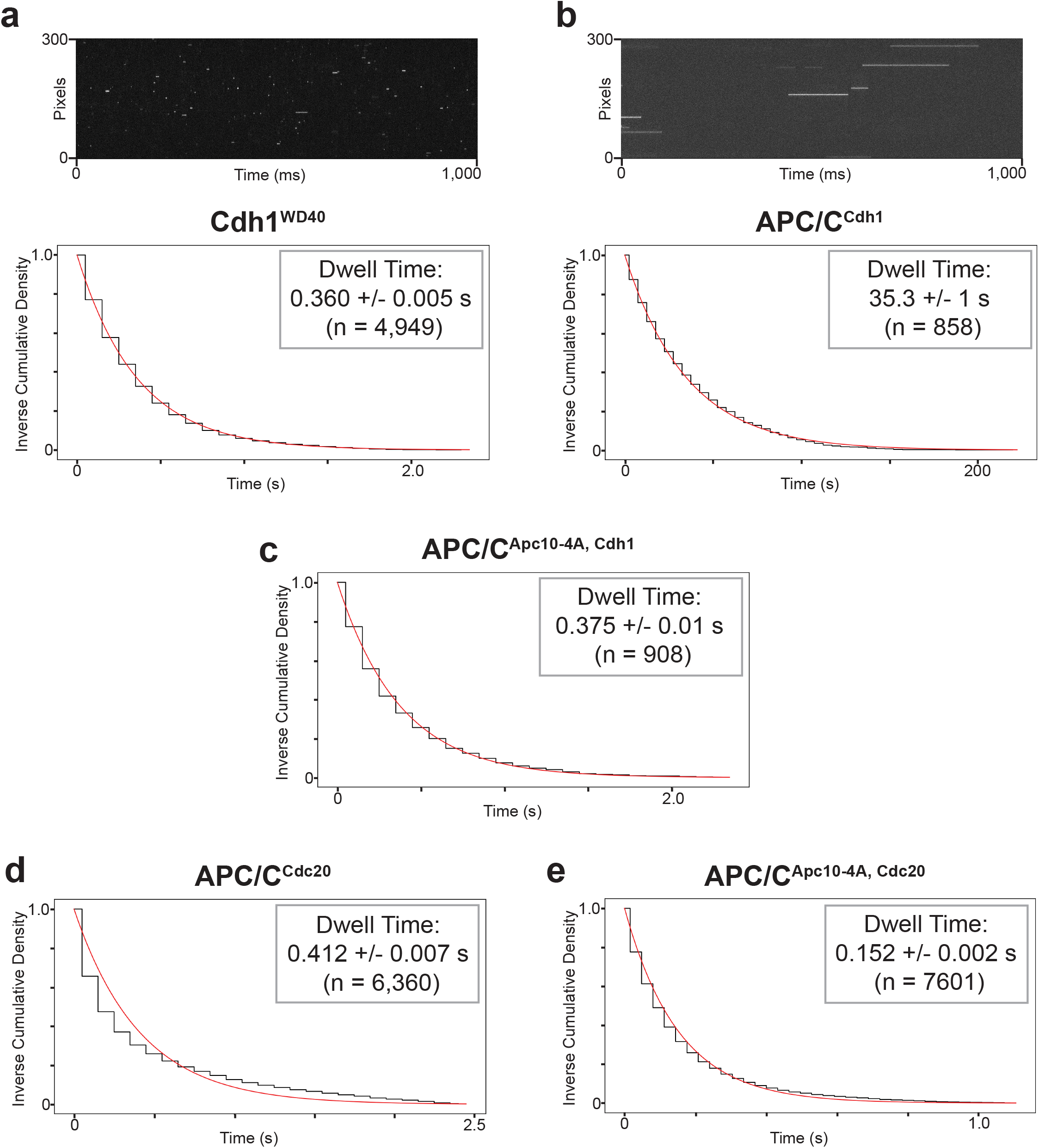
Hsl1 D box binding analysis. Dwell time distributions from SMOR analysis of representative movies with Cy5-labeled Hsl1 D box peptide and GFP-tagged binding protein on the glass surface: **a**, Cdh1^WD40^ (100 pM peptide); **b**, APC/C^Cdh1^ (100 pM peptide); **c**, APC/C^Cdh1^ with Apc10-4A mutations (1 nM peptide); **d**, APC/C^Cdc20^ (1 nM peptide); **e**, APC/CCdc20 with Apc10-4A mutations (1 nM peptide). Panels **a** and **b** include kymographs of binding events over time. Insets indicate mean dwell time +/-SEM (n indicates number of selected peaks). Results are representative of 2 independent experiments (Supplementary Table 1).

The minimum frame rate of our camera with full use of all active pixels was 32 ms. Ideally, the calculated mean dwell time should be greater than 3 times the frame rate, and thus we were unable to reliably measure dwell times less than 100 ms. There were multiple instances in which we observed single-frame interactions. Although a dwell time could not be calculated in these cases, they are likely to represent real binding and are noted in our analysis as ‘single-frame’ events.

We performed multiple independent experiments for each ligand-protein combination. A single representative replicate for each condition is described in the following sections. Additional replicates are listed in Supplementary Table 1.

### Cooperation between activator and Apc10 in D box binding

We first quantified the binding of the Hsl1 D box peptide to Cdh1 and to APC/C-activator complexes. The mean dwell time for the peptide with the Cdh1 WD40 domain was 0.360 +/- 0.005 s (Fig. 4a, Table 1; note that the error in these analyses is an estimated standard error of the mean for an exponential distribution). The dissociation rate constant k_off_ for this interaction is therefore 2.8 s^-1^. Based on the K_D_ of 4 x 10^-6^ M that we determined earlier (Fig. 1c), we infer an association rate constant k_on_ of 7 x 10^5^ M^-1^s^-1^, which is within the normal range of diffusion-limited binding events^35^.

**Table 1:**
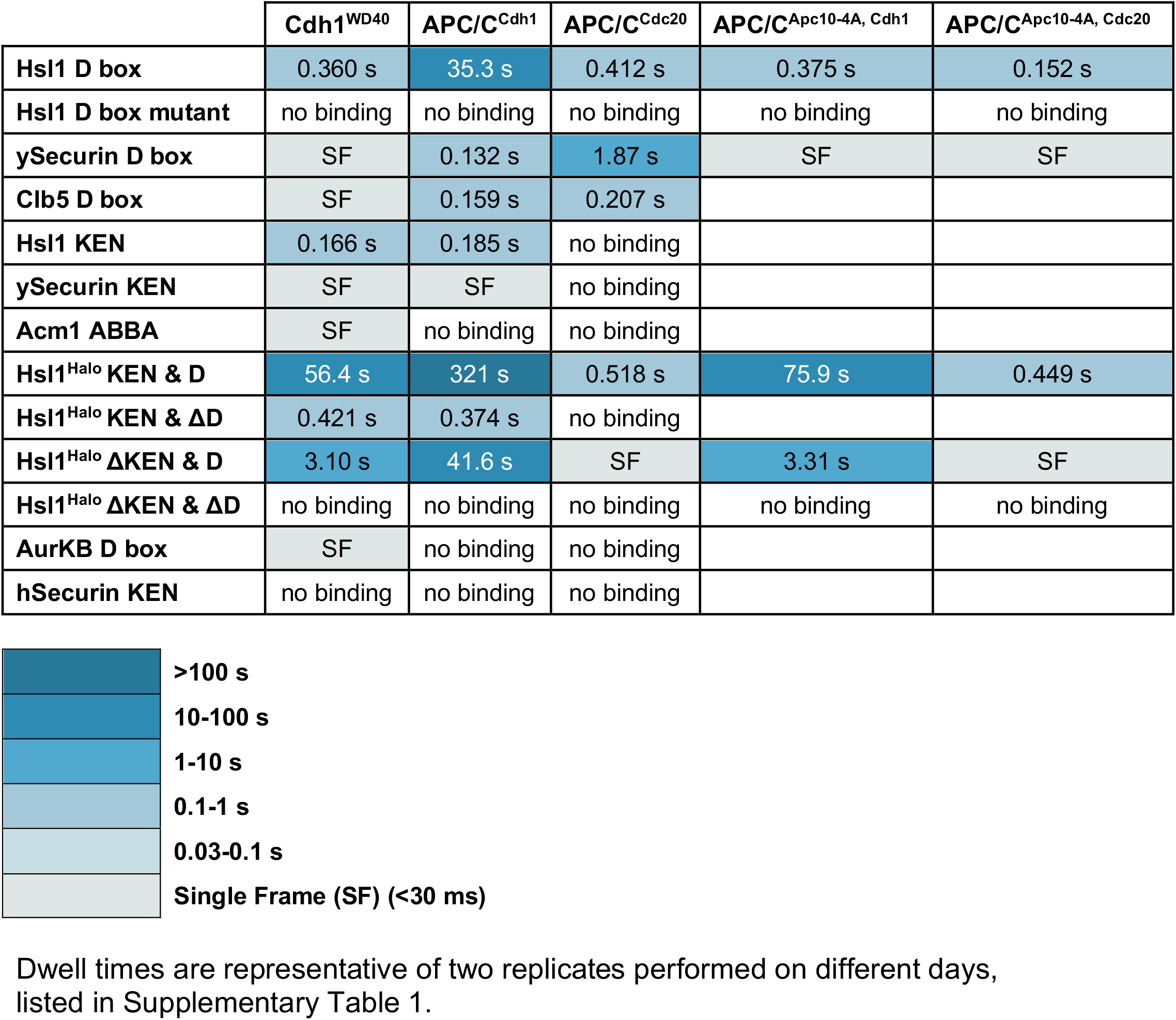
Dwell times for substrate interactions with yeast APC/C.

The dwell time of Hsl1 peptide binding to Cdh1-activated APC/C was 35.3 +/-1 s (Fig. 4b, Table 1). This ∼100-fold increase in affinity relative to Cdh1 alone seemed likely to be due to the presence of the Apc10 subunit of the APC/C, which interacts with the C-terminal end of the D box (Fig. 1a). We tested this possibility with purified APC/C containing a mutant Apc10 subunit, Apc10-4A, that contains four point mutations that eliminate D-box binding^30^. As predicted, these mutations resulted in a ∼100-fold decrease in mean dwell time to 0.375 +/- 0.01 s, which is roughly equal to the dwell time with Cdh1 alone (Fig. 4c, Table 1).

The affinity of the D box for Apc10 is known to be extremely low, as confirmed by the lack of detectable D-box binding to APC/C^apo^ (Fig 2e). Moreover, we did not observe detectable binding of the Hsl1 D box to 144 µM purified Apc10 in anisotropy experiments (Supplementary Data Fig. 5a). We conclude that a weak interaction with Apc10 cooperates with Cdh1 to provide high-affinity D-box binding. If we assume that k_on_ for D-box binding to APC/C^Cdh1^ is the same as that for binding to Cdh1 alone, then we would estimate a K_D_ of 40 nM for the binding of the Hsl1 D box to APC/C^Cdh1^.

To confirm that the APC/C^apo^ in these experiments was functional, we also tested a Cy5-labeled peptide of the C-terminal IR motif of Cdh1, which binds to the Cdc27 subunit^4^. We recovered specific protein-protein interactions with a dwell time of 0.616 +/- 0.02 s (Supplementary Data Fig. 5b, c).

The Hsl1 D box bound Cdc20-activated APC/C with a mean dwell time of 0.412 +/- 0.007 s (Fig. 4d), 85-fold lower affinity than that for APC/C^Cdh1^. Thus, the Hsl1 D box has a clear preference for Cdh1, which is consistent with evidence in vivo that Hsl1 is primarily a Cdh1 target late in mitosis. The Apc10-4A mutation reduced dwell time to 0.152 +/- 0.002 s (Fig. 4e). This 3-fold drop in dwell time is far less dramatic than the 100-fold decrease seen with APC/C^Cdh1^, perhaps suggesting that activator influences the ability of the D box to interact with Apc10; that is, the Hsl1 D box peptide does not engage with the Apc10 subunit in the same way when bound to Cdc20.

### Activator specificity of degrons

We next used the SMOR assay to analyze the other Cy5-labeled degron peptides used in our anisotropy studies (Fig 1b). In contrast to the D box of Hsl1, the ySecurin D box displayed specificity for APC/C^Cdc20^. Mean dwell time with APC/C^Cdc20^ was 1.87 +/- 0.04 s, compared with a mean dwell time with APC/C^Cdh1^ of 0.132 +/- 0.002 s (Fig. 5a). Despite being one of the earliest APC/C^Cdc20^ substrates in vivo, we found that the D box of Clb5 had similar affinity for APC/C^Cdc20^ (0.207 +/- 0.008 s) and APC/C^Cdh1^ (0.159 +/- 0.004 s) (Fig. 5b). We suspect that the early degradation of Clb5 relative to securin depends not on D box selectivity but on the presence of a Cdc20-specific ABBA motif in Clb5^15^.

**Fig. 5:**
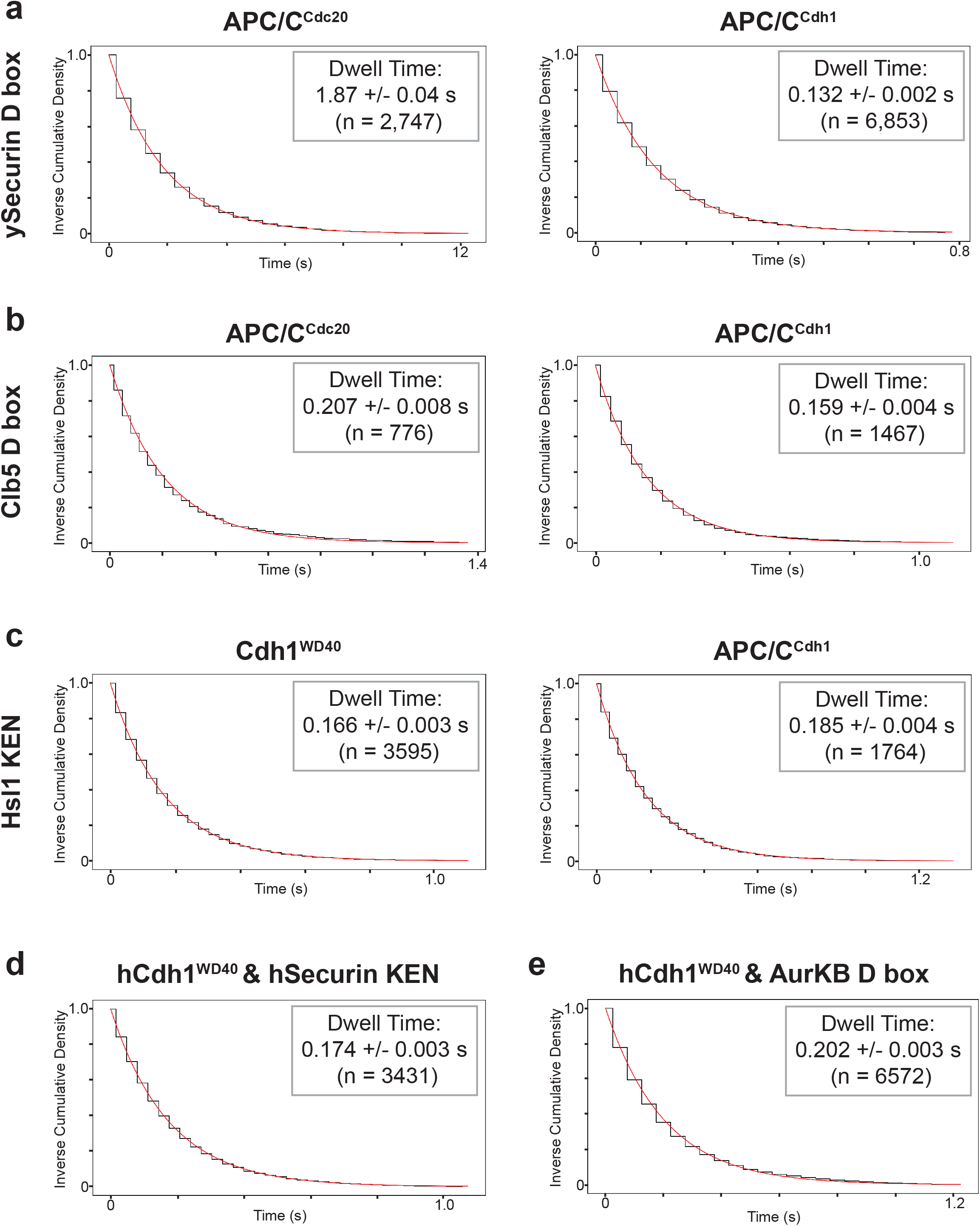
Analysis of degron from multiple substrates. **a-c**, Dwell time distributions from SMOR analysis of representative movies with Cy5-labeled ySecurin D box peptide (**a**, 1 nM), Clb5 D box peptide (**b**, 1 nM), and Hsl1 KEN peptide (**c**, 1 nM) binding to GFP-tagged Cdh1^WD40^ (left) or APC/C^Cdh1^ (right). **d**, **e**, Analysis of hSecurin KEN box peptide (**d**, 500 pM) or AurKB D box peptide (**e**, 1 nM) binding to GFP-tagged human Cdh1 WD40 domain. Insets indicate mean dwell time +/- SEM (n indicates number of selected peaks). Results are representative of 2 independent experiments (Supplementary Table 1).

The KEN peptide from Hsl1 bound Cdh1 and APC/C^Cdh1^ with similar affinity (Fig. 5c), consistent with the idea that this degron binds to the activator and not to other APC/C subunits. The KEN peptide from ySecurin bound only transiently to Cdh1 (single-frame events), suggesting a low affinity. There was no detectable binding of either KEN peptide to APC/C^Cdc20^ (Table 1).

We also analyzed interactions between human activators and degrons. We were able to prepare bulk quantities of the WD40 domains of Cdc20 and Cdh1 for fluorescence anisotropy studies. We observed good binding (K_D_ ∼ 2 µM) to both activators by the Hsl1 D box and significant but low-affinity binding to Cdh1, but not Cdc20, by the KEN box from human securin/Pttg1 (hSecurin) and D box from Aurora B kinase (AurKB) (Supplementary Data Fig. 6). We also used single-molecule studies to analyze the binding of various degrons to human activators (Table 2). The hSecurin KEN peptide did not bind human Cdc20^WD40^ but bound human Cdh1^WD40^ with a dwell time of 0.174 +/- 0.003 s (Fig. 5d, Supplementary Data Fig. 7, Table 2). Similarly, the AurKB D box bound human Cdh1^WD40^ with a dwell time of 0.202 +/- 0.003 s (Fig. 5e, Supplementary Data Fig. 7) but displayed lower affinity for Cdc20 (single-frame interactions only).

**Table 2:** Dwell times for substrate interactions with human activators.

|  | **hCdh1<sup>WD40</sup>** | **hCdc20<sup>WD40</sup>** |
| --- | --- | --- |
| **Hsl1 D box** | 4.52 s | 1.08 s |
| **Hsl1 D box mutant** | no binding | no binding |
| **ySecurin D box** | SF | SF |
| **Clb5 D box** | SF | 0.054* s |
| **Hsl1 KEN** | 1.48 s | SF |
| **ySecurin KEN** | 0.869 s | SF |
| **Acm1 ABBA** | no binding | no binding |
| **AurKB D box** | 0.202 s | SF |
| **hSecurin KEN** | 0.174 s | no binding |

As in Table 1, dwell times are representative of two replicates performed on different days, listed in Supplementary Table 1. Colors as in Table 1.
\*Calculated dwell time is less than 3 times the frame rate and is therefore less reliable.

### Substrate with multiple degrons binds with very high affinity

APC/C substrates often contain multiple degrons. To fully understand substrate interactions with the APC/C, we therefore analyzed its interaction with a substrate carrying both D box and KEN degrons. We used a well-studied fragment of Hsl1 (amino acids 667-872)^29^, tagged with a C-terminal HaloTag to which chemical dye JF549 covalently binds^36^ (Fig. 6a). APC/C ubiquitylation assays and an APC/C ensemble binding assay confirmed that the HaloTag does not affect ubiquitylation or binding (Supplementary Data Fig. 2d, e). This substrate, including mutants lacking one or both degrons, was tested under the same conditions as those in our peptide binding experiments and found to have specific single molecule binding (Supplementary Data Fig. 8). The dwell times are summarized in Table 1.

**Fig. 6:**
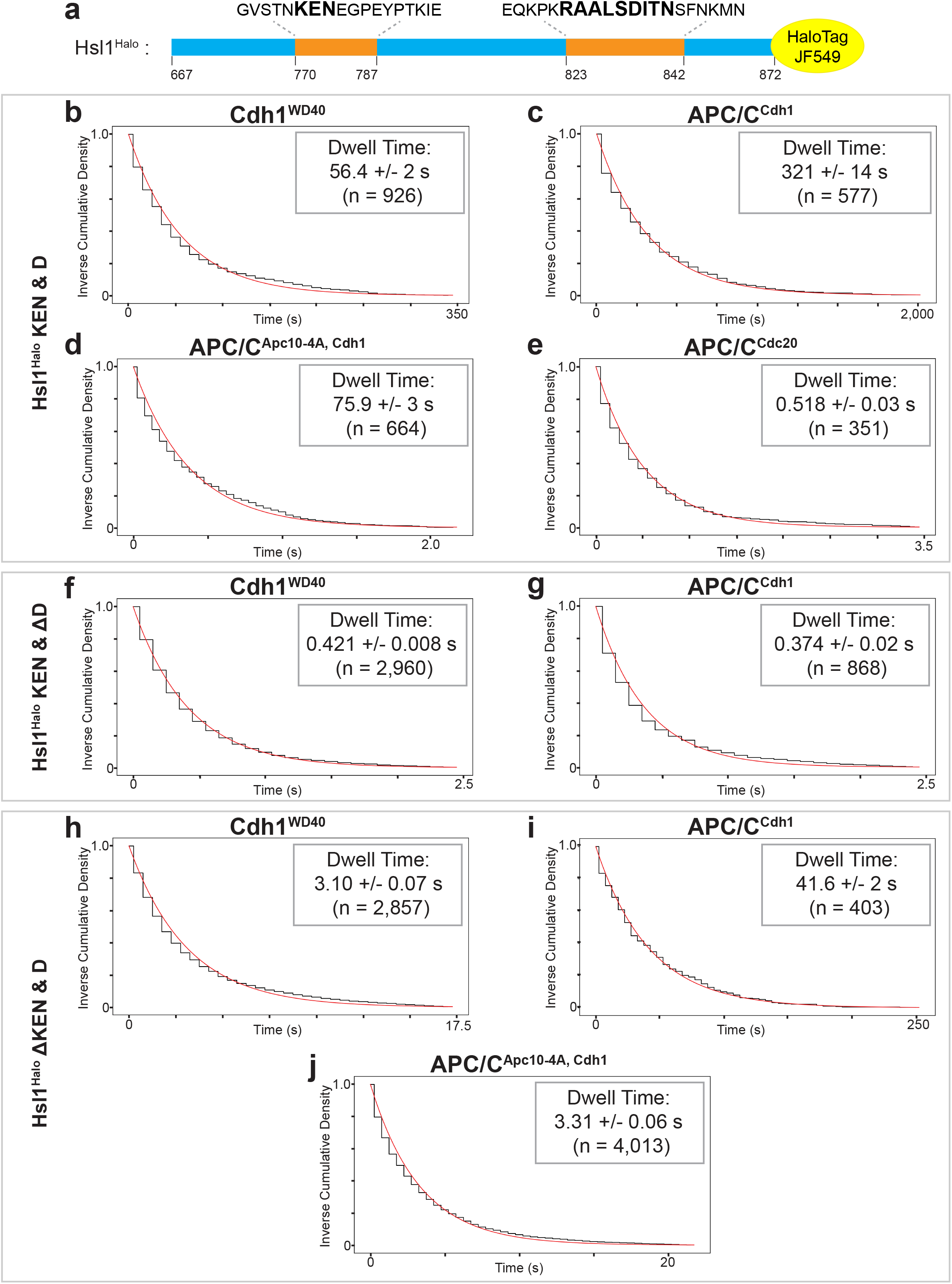
Double degron substrate interactions. **a**, Hsl1^Halo^ fragment used in these experiments, showing sequences of KEN and D boxes. **b-e**, Dwell time distributions from SMOR analysis of representative movies with Hsl1^Halo^ carrying wild type degrons and the indicated GFP-tagged binding proteins: **b**, Cdh1^WD40^; **c**, APC/C^Cdh1^; **d**, APC/C^Cdh1^ with Apc10-4A mutations; **e**, APC/C^Cdc20^. **f-g**, Analysis of Hsl1^Halo^ with mutant D box, binding to: **f**, Cdh1^WD40^; **g**, APC/C^Cdh1^. **h-j**, Analysis of Hsl1^Halo^ with mutant KEN, binding to: **h**, Cdh1^WD40^; **i**, APC/C^Cdh1^; **j**, APC/C^Cdh1^ with Apc10-4A mutations. Insets indicate mean dwell time +/- SEM (n indicates number of selected peaks). Results are representative of 2 independent experiments (Supplementary Table 1).

The combination of both a KEN and D box in a single substrate resulted in extremely high affinity binding. The dwell time with Cdh1^WD40^ increased 100-fold from ∼0.4 s and ∼0.2 s for D and KEN box peptides, respectively, to 56.4 +/- 2 s for the Hsl1 fragment (Fig. 6b). The interaction was increased another 6-fold with APC/C^Cdh1^ (dwell time 321 +/- 14 s; Fig. 6c). This boost in affinity was not seen in the Apc10-4A mutant (Fig. 6d), as in our earlier studies of the D box alone (Fig. 4c). Again, applying the k_on_ calculated from anisotropy experiments, we infer that the dissociation constant of Hsl1 for APC/C^Cdh1^ is ∼4 nM.

Activator specificity was retained by the Hsl1 fragment, as the dwell time of 0.518 +/- 0.03 s with APC/C^Cdc20^ is 600-fold lower than that with APC/C^Cdh1^ (Fig. 6e). This specificity can also be seen in the processivity of ubiquitylation of this substrate (Supplementary Data Fig. 2f). Interestingly, as seen in our studies of the Hsl1 D box peptide, the Apc10-4A mutations only slightly reduced Hsl1 dwell time with APC/C^Cdc20^, suggesting as before that the Hsl1 D box does not engage effectively with Apc10 when bound to Cdc20.

An Hsl1 fragment carrying mutations in the D box was also tested. We would expect this substrate to interact primarily through the KEN box. This substrate displayed a recoverable dwell time with Cdh1^WD40^ at 0.421 +/- 0.008 s and APC/C^Cdh1^ at 0.374 +/- 0.02 s, but not with APC/C^Cdc20^ (Fig. 6f, g and Table 1). These dwell times are very similar to those observed with the KEN peptide, further suggesting that the KEN motif binds poorly to Cdc20.

We also tested an Hsl1 fragment with mutations in the KEN box. This substrate was expected to bind primarily through the D box. Interestingly, we found a dwell time of 3.10 +/- 0.07 s with Cdh1^WD40^, a 10-fold increase from that with the D box peptide alone (Fig. 6h, Table 1). The interaction of this mutant substrate with APC/C^Cdh1^ was similar to that with the D box peptide (Fig. 6i). To test if the 10- fold increase with Cdh1 alone was due to Hsl1^Halo^ interacting with a portion of the WD40 domain that is occluded by the APC/C, we also tested this mutant with Cdh1-activated APC/C^Apc10–4A^. This interaction occurred with a dwell time of 3.31 +/- 0.06 s, which is similar to that with Cdh1^WD40^ (Fig. 6j). These results suggest that an Hsl1 sequence outside the tested D box peptide interacts with Cdh1 in a way that is blocked by the interaction with Apc10.

## Discussion

Quantitative analysis of macromolecular interactions is often hindered by the low protein yields that result from heterologous overexpression and purification. We addressed this problem by developing a straightforward, adaptable, and robust single-molecule binding assay that requires minimal amounts of protein (nanograms). We used the SMOR method to carry out quantitative analyses of APC/C interactions with its substrates, providing new insights into the specificity of substrate binding for different activators, and the role of multivalent degron interactions in high-affinity binding.

The activators of the APC/C are thought to possess distinct substrate specificities^15, 21^: Cdc20 triggers anaphase by promoting degradation of a small number of substrates (including securin and Clb5), while in late mitosis and G1 Cdh1 promotes degradation of an expanded range of substrates (including Hsl1). Ubiquitylation of securin and Clb5 by APC/C^Cdc20^ is more processive than that with APC/C^Cdh1^, while Hsl1 is more processively modified by APC/C^Cdh1^ (ref ^19^). We now provide direct quantitative evidence to demonstrate activator specificity in degron binding. We find that the yeast securin D box displays 15-fold higher affinity for APC/C^Cdc20^, while the Hsl1 D box has 85-fold preference for APC/C^Cdh1^. These preferences presumably depend on residues in the degron other than the conserved RxxL consensus, which interact with specific features of the binding site on the activator.

Differences in the D-box binding site of the activators is further supported by the Cdc20 specificity of the chemical inhibitor Apcin^37^. This specificity raises the exciting possibility of activator-specific targeted protein degradation as a therapeutic application.

Surprisingly, the D box of Clb5 displays similar (moderate) affinity for both activators despite its preference for Cdc20 in vivo and in ubiquitylation assays. It seems likely that Cdc20 specificity in the case of Clb5 is provided by its ABBA motif, which is known to be required for early Clb5 degradation and is specific for yeast Cdc20^15^. Thus, activator specificity depends on the D box in some cases while in others is provided by a second degron.

Interestingly, the KEN box of securin is specific for APC/C^Cdh1^ and does not bind APC/C^Cdc20^. Residues outside the KEN motif must influence binding to different features on the two activators. As securin is known to be preferred by Cdc20 in vivo and in ubiquitylation assays, the preference of its KEN box for Cdh1 must not overcome the stronger preference of its D box for Cdc20.

We did not observe any binding of KEN peptides to Cdc20 using yeast or human Cdc20 with yeast or human KEN peptides, respectively (Tables 1, 2). The only case in which we observed an interaction was single-frame binding of yeast KEN peptides to human Cdc20 (Table 2). The KEN degron was originally identified as a motif targeted by Cdh1^12^, and there is evidence to support Cdh1 specificity of the KEN box in some substrates. In Cyclin A2, for example, the KEN box is more important for ubiquitylation by APC/C^Cdh1^ than by APC/C^Cdc20^, whereas D boxes are more important for ubiquitylation by APC/C^Cdc20^ (ref ^18^). KEN degrons might increase the Cdh1 affinity of substrates with Cdc20-specific D boxes, ensuring that these substrates continue to be unstable in late mitosis and G1.

In contrast to our evidence that the KEN box binds poorly, if at all, to Cdc20, there is structural evidence for Cdc20 binding to KEN degrons from BubR1 and Cyclin A2^18, 38^. In these structures the KEN box is part of a protein containing additional degrons, and it seems likely that KEN binding to its low-affinity site on Cdc20 is driven in these cases by the high local concentration provided by a multivalent ligand.

Our studies also provide a quantitative understanding of the contributions of activator and Apc10 to the composite D-box binding site. The affinity of the Hsl1 D box for APC/C^Cdh1^ is 100-fold higher than that with Cdh1 alone or with an APC/C carrying an Apc10 mutation, showing the dramatic impact of Apc10 on D box affinity. D-box binding can be considered as a bivalent interaction, in which the N-terminal RxxL segment of the degron binds with moderate affinity (K_D_=4 µM) to specific sites on the activator surface, while poorly-conserved sequences at the C-terminal end of the D box interact with Apc10. The latter interaction is not well understood at the structural level and is clearly very low affinity. We observed no binding of D box peptides to APC/C^apo^ or Apc10, and the only reported evidence for direct binding comes from NMR analysis of Apc10-D box interactions at high (5 mM) concentrations of Hsl1 D box peptide^39^. Nevertheless, this low-affinity Apc10 interaction cooperates effectively with the moderate-affinity activator interaction to generate high-affinity bivalent binding of the D box to APC/C^Cdh1^.

The D-box binding pocket is well conserved in Cdc20 and Cdh1, but our results suggest that the two activators present the D box to Apc10 in different ways. Although Apc10 boosted affinity for the Hsl1 D box by 100-fold in the case of APC/C^Cdh1^, it seemed to provide only a ∼3-fold increase in binding to APC/C^Cdc20^. Similarly, binding to APC/C^Cdc20^ of Hsl1 containing both D and KEN boxes is only slightly reduced by mutation of Apc10. Perhaps the Cdc20-D box complex is oriented in a way that results in a low-affinity interaction with Apc10^18, 40^.

Most if not all APC/C substrates contain multiple degrons, and our studies document the high affinity that results from the multivalent binding of D and KEN boxes of Hsl1. When a moderate affinity D box (K_D_∼4 µM) exists on the same protein as a moderate affinity KEN box (K_D_∼12 µM), the result is an Hsl1 dwell time of 300 seconds – suggesting a dissociation constant of ∼4 nM. The effects of multivalency are further enhanced by allosteric enhancement of each degron’s binding when the other is bound (Fig. 1d).

The Hsl1 dwell time of 5 min is a very long time in the life of a yeast cell, which divides every 90 minutes. This raises the possibility that Hsl1 does not dissociate spontaneously from the APC/C but is extracted from the APC/C by the proteasome. However, Hsl1 may be an unusual case, as suggested by the unusually high affinity of its degrons. The degrons of securin and Clb5 have lower affinities than those of Hsl1, and these substrates are therefore likely to bind with lower affinity. Unfortunately, we were unable to measure the binding of securin and other substrates due to their tendency to aggregate and create excessive background fluorescence in our assay.

In sum, our results reveal that the affinity and specificity of the APC/C-substrate interaction can be influenced by a remarkable array of factors. Key factors include the specificity and affinity of individual degrons for the activator and Apc10, as well as the presence of multiple degrons on a substrate. Numerous other factors are also likely to be important, such as the number and positioning of lysines for modification, the distance between degrons, and the orientation of the D box at its bivalent binding site^7,16, 18, 40^.

The timing of substrate ubiquitylation and destruction is important for robust control of cell cycle events. Our past work suggests that substrate affinity is a key determinant of the timing of substrate degradation^15, 22^, and there might be some contribution from competition among substrates^23^. A full understanding of the ordering of substrate degradation will require more extensive studies of the concentrations and affinities of substrates and the APC/C inside the cell.

Using tools developed by single-molecule biophysics, the SMOR assay provides a straightforward approach for biologists and biochemists to study macromolecular interactions that are difficult to study by conventional methods. Although our experiments were performed with purified components, we suspect that the SMOR assay will also be effective for studying binding proteins that are purified directly on the glass^31^. The continued development of single-molecule approaches promises to open many new avenues in the study of biological and therapeutic interactions.

## Methods

### Yeast APC/C purification

Yeast strains were derivatives of W303 and are listed in Supplementary Table 2. For APC/C purification, we used a strain carrying Cdc16-TAP and lacking Cdh1 (DOM1126); in most experiments the strain also carried Apc1-GFP (NHY13). Yeast were grown in YPD media to OD_600_ = 0.8, collected and flash frozen. Cells were lysed by bead beating in lysis buffer A (50 mM HEPES pH 7.5, 150 mM KCl, 10% glycerol, 0.2% Triton X-100, 63 µM B-glycerophosphate, 48 µM sodium fluoride, 1 µg/ml pepstatin A, 1 µg/ml leupeptin, 1 µg/ml aprotinin, 1 mM PMSF, 1 mM DTT, 0.5 mM EDTA) and APC/C was purified using magnetic IgG beads. The beads were washed using wash buffer A (20 mM HEPES pH 7.5, 150 mM KCl, 10% Glycerol, 0.05% Triton X-100). After incubation with purified Cdh1 or Cdc20, the APC/C was cleaved off the beads with TEV protease in wash buffer A with 0.05% Tween-20 (for single molecule studies) or 0.05% Triton X-100 (for ensemble assays such as ubiquitylation) and used immediately.

### Activator purification

Activators were cloned into the pFastBac HT A vector, with an N-terminal 2xStrep-Tag II. For some experiments, we constructed vectors for expression of the WD40 domains of yeast Cdh1 (aa 241 to 550; pNH144), human Cdh1 (aa 165 to 484; pNH164), or human Cdc20 (aa 162 to 484; pNH188). For SMOR, a C-terminal GFP tag was added. Bacmid plasmids are listed in Supplementary Table 3. Plasmids were transformed into DH10Bac cells, and purified Bacmid was used to transfect Sf9 cells to generate P1 baculovirus, which was used to generate P2 virus. SF9 cells were infected with P2 virus for 48 h. Flash-frozen pellets were lysed by sonication or high-pressure homogenization in lysis buffer B (50 mM Tris-HCl pH 8, 150 mM KCl, 0.5 mM EDTA, 1 mM DTT, 10% Glycerol, EDTA-free protease inhibitor tablet, 1 µg/ml pepstatin A). After the lysate was applied to the StrepTrap column, the column was washed with wash buffer B (lysis buffer B lacking protease inhibitors) and eluted in the same buffer containing 2.5 mM desthiobiotin.

### Polarization Anisotropy

Fluorescent degron peptides carried a C-terminal Cy5 label (CPC Scientific) and are listed in Supplementary Table 4. For anisotropy experiments, 10 nM fluorescent peptide was mixed with various concentrations of purified Cdh1 WD40 in wash buffer B at room temperature for 1 min, which we determined was sufficient for the binding reaction to reach equilibrium. Fluorescence was measured on a K2 Multifrequency Fluorometer at 25°C. All Cy5-labeled peptides were excited with polarized light at 635 nm and emission was detected using a 700/75 nm bandpass filter (ET series, Chroma). A competition experiment was conducted with unlabeled Hsl1 D box peptide to confirm that the Cy5 dye did not bind the Cdh1 WD40. Data were fitted to one-site equilibrium binding using Prism 8 to determine K_D_. Results were the same when peptide binding was measured in the buffer used in SMOR assays (buffer C, below).

### SMOR assay surface preparation and protein immobilization

For all SMOR experiments, 24×50 mm high precision glass cover slips (Bioscience Tools) and drilled microscope slides were passivated with a combination of PEG and PEG-biotin (cover slips) or PEG only (slides) following a previously published protocol for SiMPull^31^. Reaction chambers (∼20 µl) were created using double-sided tape and epoxy. Protein was immobilized on the surface with 0.2 mg/ml NeutrAvidin in buffer C (20 mM HEPES pH 7.5, 150 mM KCl, 10% glycerol, 0.05% Tween-20, 1 mM DTT). Excess NeutrAvidin was washed away and incubated with either biotin-conjugated mouse monoclonal anti-Strep-Tag II antibody (LifeSpan BioSciences, Inc.) or biotin-conjugated rabbit polyclonal anti-GFP antibody diluted in buffer C + BSA (0.1 mg/ml Molecular Biology Grade Bovine Serum Albumin). Activator WD40 and activated APC/C (typically 50 µl containing about 65 ng APC/C, of which only a small fraction is immobilized on the glass) were immobilized using anti-Strep-Tag II, while APC/C alone was immobilized with anti-GFP. SMOR results were similar when performed in the buffer (wash buffer B) used in polarization anisotropy.

### Kinetics Experiments with SMOR assay

To capture dynamic protein-protein interactions, dye-labeled substrate diluted in buffer C + BSA was added to the chamber containing immobilized proteins. Interactions were imaged by TIRF microscopy as described below. For optimal signal-to-noise ratio, peptide concentration was no greater than 1 nM. Protein substrates included Hsl1 aa 667-872 and a C-terminal HaloTag followed by a TAP tag, and were produced by translation in rabbit reticulocyte lysates using TnT Quick Coupled Transcription/Translation System (Promega) (see Supplementary Table 5 for plasmids). Protein was purified with magnetic IgG beads and labeled on the bead with JF549 or JF646 dye so that unbound dye could be washed away. Purified, dye-labeled protein was then cleaved from the beads using TEV protease. Before adding substrate to the reaction chamber, oxygen scavenging reagents were added (10 nM protocatechuate-3,4- dioxygenase, 2.5 mM protocatechuic acid, and 1 mM Trolox)^41^. Detailed description of the SMOR analysis pipeline is found in Supplementary Methods.

### Single Molecule TIRF microscopy

Microscopy was performed with a Nikon Eclipse TE2000-E with Perfect Focus with a 100×1.49na oil, Apo TIRF DIC N2, 0.13-0.2, WD0.12 objective. Movies were recorded using the Andor iXon DU-897E-CSO. For visualizing GFP-tagged protein, a 491 nm laser was used with an ET525/50 (Chroma) filter for emission. For visualizing JF549-labeled substrate, a 561 nm laser was used with an ET595/50 (Chroma) filter for emission. For visualizing all Cy5-labeled peptides and JF646-labeled substrates, a 640 nm laser was used with an ET685/70 filter (Chroma) for emission. Initially, for each substrate tested, we acquired movies at multiple intervals to deduce the optimal interval to decrease bleaching. µManager was used to control the microscope and record time-lapse movies.

### Ubiquitylation Assay

APC/C ubiquitylation assays were performed as described previously^24^. In short, APC/C was purified from yeast as described above using magnetic IgG beads and activated with purified Cdh1 or Cdc20 activator. For all APC/C ubiquitylation assays, substrates were produced by in vitro translation with ^35^S-Methionine (See Supplementary Table 5 for plasmids). Substrates were purified using magnetic IgG beads. E1 and E2 (Ubc4) were expressed in *E. coli* and purified as described previously^25, 42^. E2 was charged at 37°C for 30 min (for yeast: reaction contained mg/ml Uba1, 2 mg/ml Ubc4, 2 mg/ml methylated ubiquitin from Boston Biochem #Y-501, and 1 mM ATP; for human: same as yeast but 10 μM UbcH10 was used for E2). Charged E2 was added to a reaction containing activated APC/C and substrate. Reaction products were analyzed by SDS-PAGE and Phosphorimaging on a Typhoon 9400 Imager and quantified using ImageQuant (GE Healthcare).

## Acknowledgements

We thank Nico Stuurman, DeLaine Larson, and Kurt Thorn for their expert advice on microscopy, Ron Vale for advice and support, Luke Lavis and Margaret Jefferies (HHMI) for generously sharing fluorescent dyes, Diego Garrido Ruiz and Matt Jacobson for discussions of activator structure, and Sy Redding, Lucy D. Brennan, and Stephanie L. Johnson for their invaluable advice about single-molecule analysis. We would also like to thank members of the Morgan lab for valuable discussions and Hayk R. Mangasarian for his patience. This work was supported by a Graduate Research Fellowship from the National Science Foundation, the Paul and Daisy Soros Fellowship for New Americans, the UCSF Discovery Fellowship, and the National Institute of General Medical Sciences (R35-GM118053, to D.O.M.).

## Contributions

N.H. and D.O.M. conceived the single-molecule methods and experiments. N.H. designed and performed the experiments. J.S. created the image analysis code. A.J. helped develop single-molecule methods. N.H. and D.O.M. wrote the paper with assistance from all authors.

## Supplementary Material

**Supplementary Data Fig. 1:**
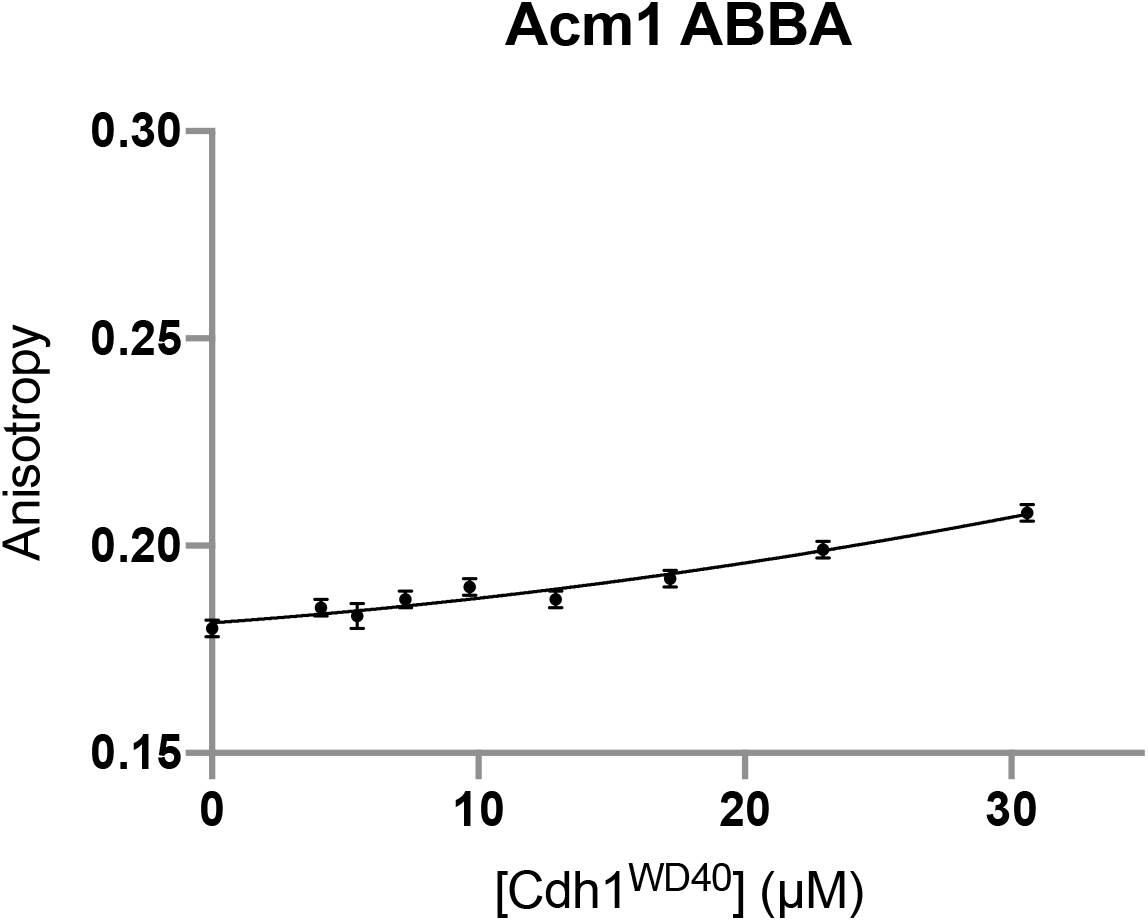
Anisotropy with Acm1 ABBA degron peptide. Fluorescence anisotropy analysis of Cy5-labeled Acm1 ABBA degron peptide (SKAAQFMLYEETAEERNI-K[Cy5]; 10 nM) binding to Cdh1^WD40^. Data points represent mean +/- SD (n = 10 reads per reaction).

**Supplementary Data Fig. 2:**
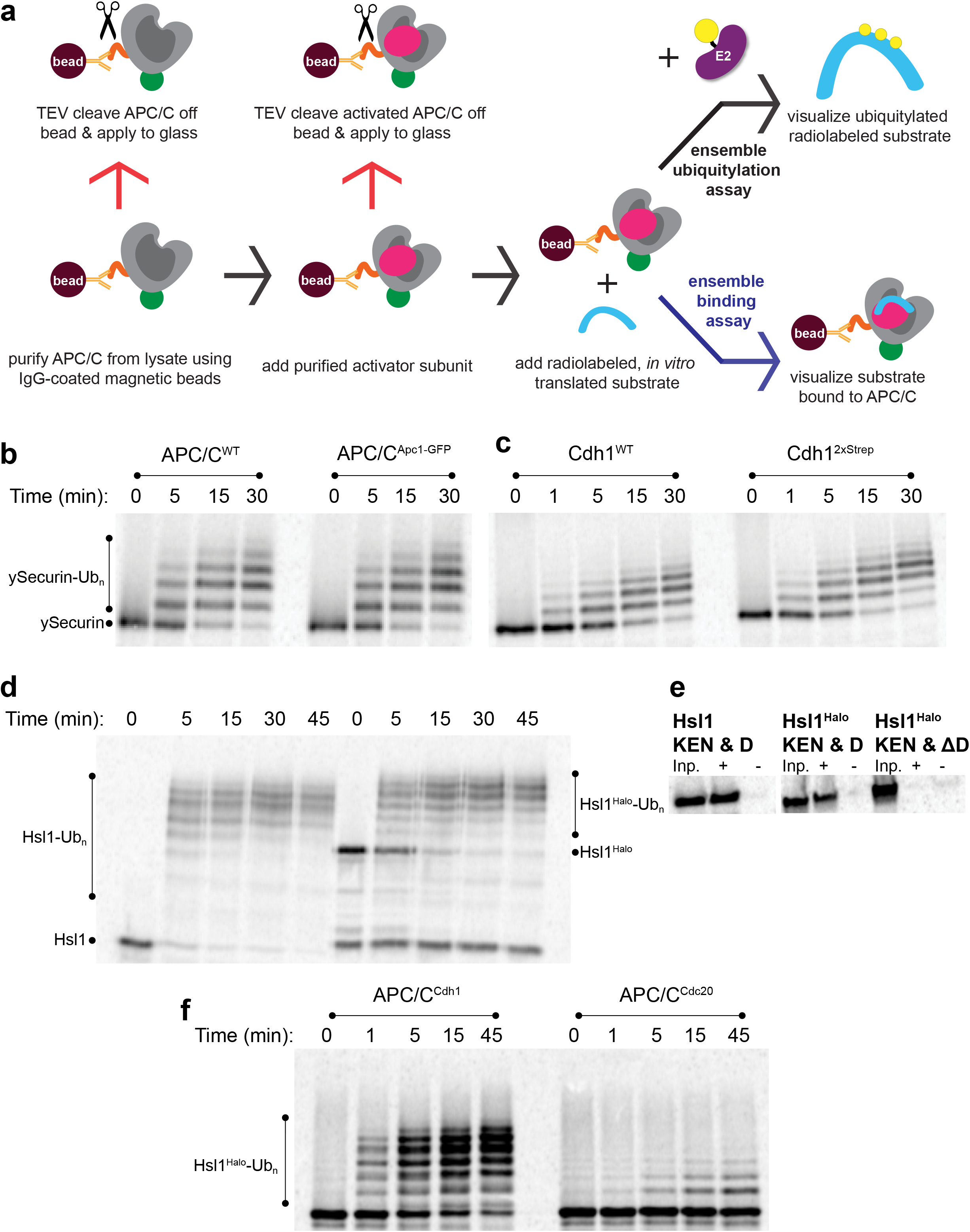
APC/C activity with tagged substrates. **a**, Methods for analysis of APC/C activity and substrate binding. **b**, APC/C ubiquitylation assay with radiolabeled full-length ySecurin, comparing wild-type APC/C^Cdh1^ and APC/C^Cdh1^ with a GFP tag on Apc1. **c**, Comparison of activities with APC/C^Cdh1^, using either wild-type Cdh1 or Cdh1 with the N-terminal 2XStrep-Tag II. **d**, Comparison of activities with APC/C^Cdh1^ and the Hsl1 fragment (667- 872) with and without the C-terminal Halo tag. **e**, Binding of radiolabeled Hsl1 fragments to yeast APC/C^Cdh1^ immobilized on magnetic beads. First lane (Inp, input) indicates the amount of labeled protein added to the beads prior to washing, second lane (+) indicates amount bound, and third lane (-) indicates background binding in absence of APC/C^Cdh1^. **f**, Ubiquitylation of Halo-tagged Hsl1 fragment with APC/C^Cdh1^ and APC/C^Cdc20^.

**Supplementary Data Fig. 3:**
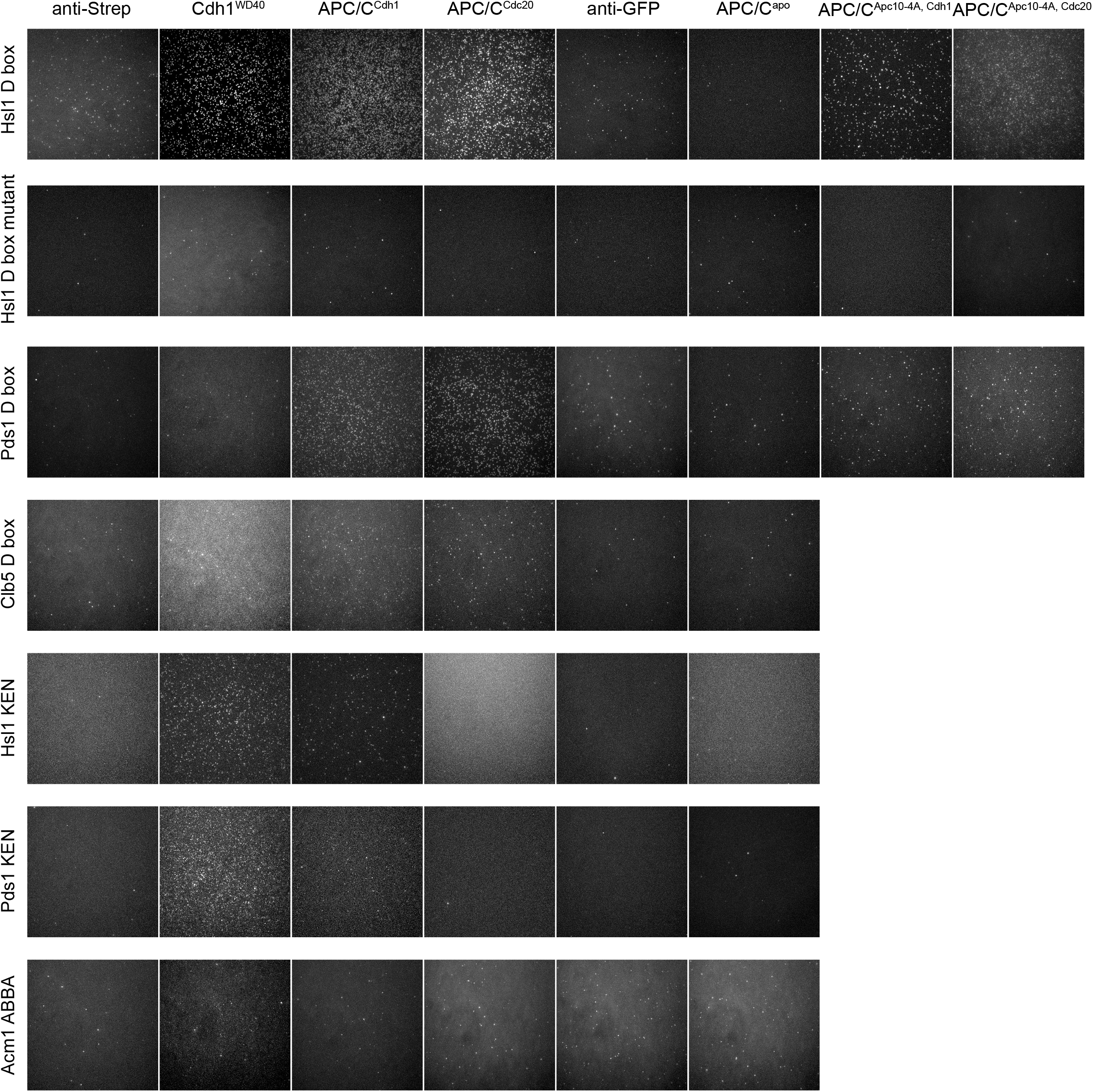
Yeast degron peptide binding in various conditions. Maximum intensity projections of 500 frames from videos of Cy5-labeled degron peptides (labeled on left) binding to various immobilized binding proteins (labeled at top). Images labeled ‘anti-Strep’ are background controls for binding to APC/C-activator complexes; images labeled ‘anti-GFP’ are background controls for binding to GFP-tagged APC/C^apo^ and Cdh1^WD40^.

**Supplementary Data Fig. 4:**
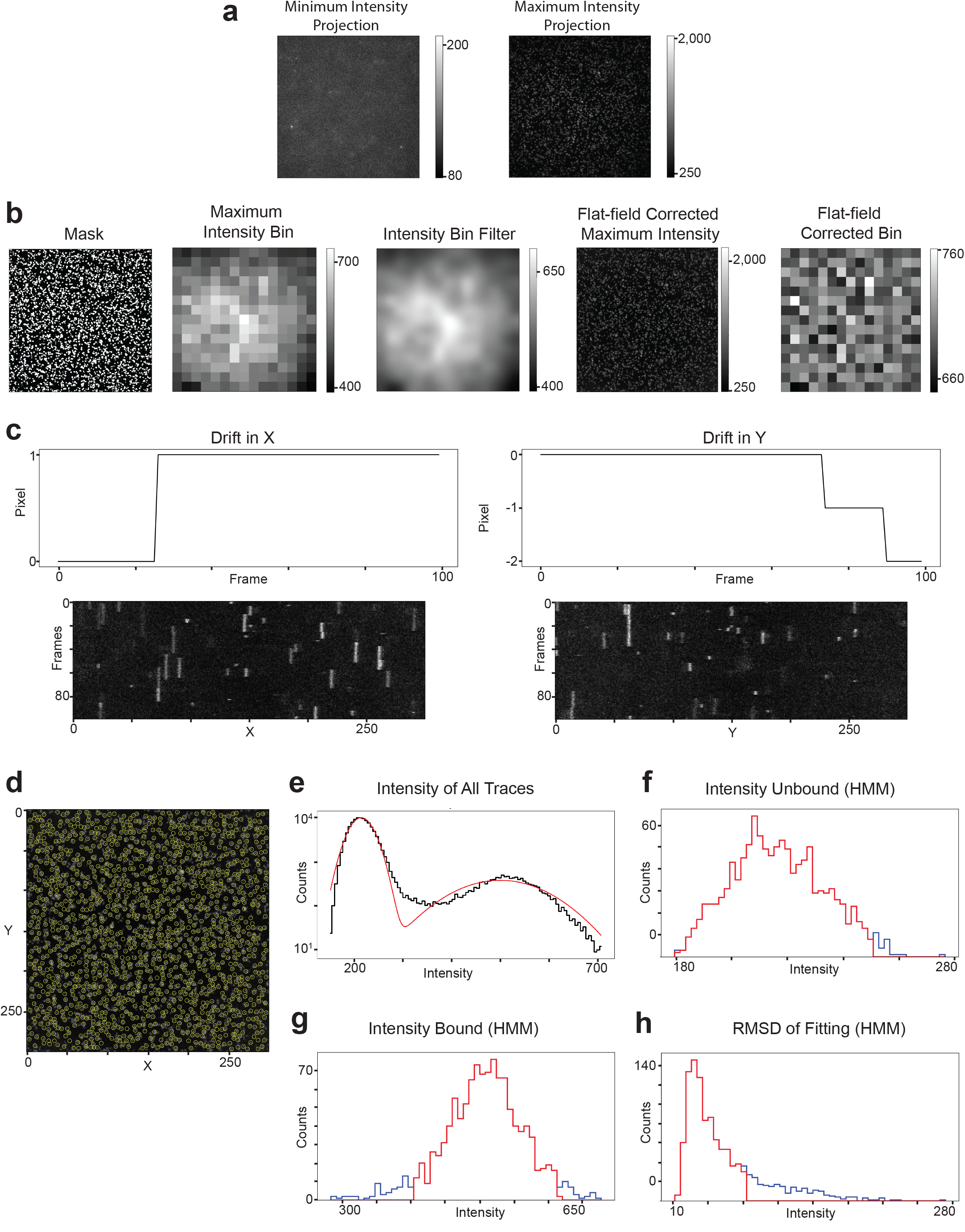
Steps in the SMOR analysis process. **a**, As a first step in the analysis, the code produces figures for maximum and minimum intensity projections (Z-stack) of the movie being analyzed. **b**, A mask is created by a binary filter to find the pixels with bright spots. The mean intensity within the mask is calculated for each 20×20 pixel bin, and a Gaussian filter is applied to create the intensity bin filter. Flat-field correction is performed by normalizing the maximum intensity projection by the filter. As described in the main text, the process of flatfield correction produces several figures for the user to track the process (see Fig. 3b). **c**, Drift correction produces a plot to show the number of pixels of drift in both x and y axes as well as a kymograph in both x and y of the corrected movie for the user to determine if the drift has been adequately corrected. Each binding event produces a straight line in the kymograph, and drift results in a 1-2-pixel shift in all lines. **d**, The code initially identifies (x,y) coordinates where binding occurs in the movie by using a peak-finding algorithm and creates this peak identification plot. **e**, A histogram of minimum and maximum intensities along the entire trace (in time or z) at each (x,y) coordinate where a binding event was identified. Red line indicates double Gaussian fit of the data. **f**, Histogram of unbound intensities after HMM fitting. **g**, Histogram of bound intensities after HMM fitting. **h,** Plot of root mean squared deviation (RMSD) between the experimental trace and the HMM trace fitting among all (x,y) coordinates. For panels **f**-**h**, red indicates selected data (≤2 SD from the median) and blue indicates rejected data.

**Supplementary Data Fig. 5:**
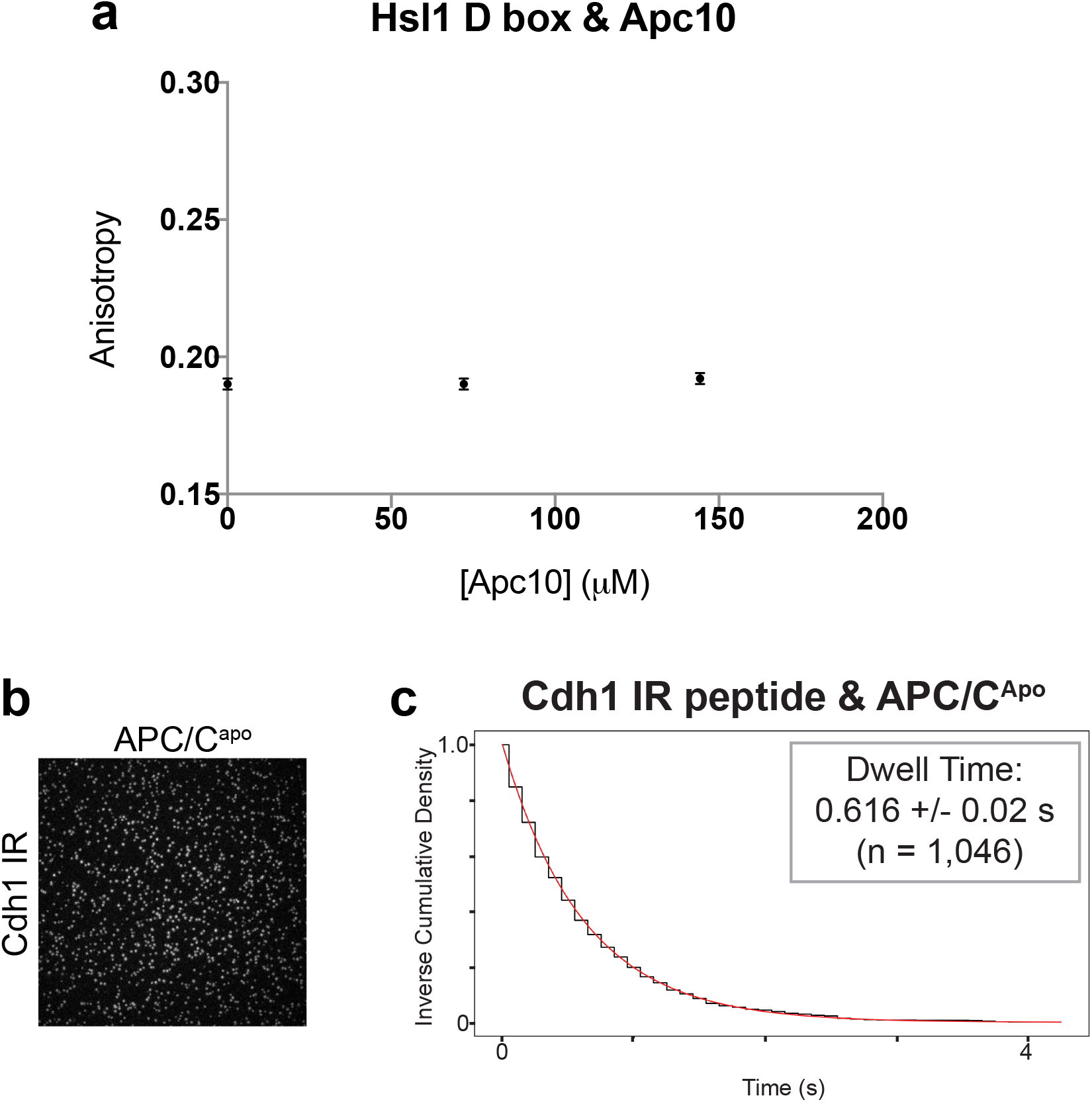
Control experiments with Apc10 and APC/C^apo^. **a**, Fluorescence anisotropy analysis of 10 nM Cy-5 labeled Hsl1 D box peptide incubated with up to 144.3 µM purified yeast Apc10. Data points represent mean +/- SD (n = 10 reads per reaction). **b**, Maximum intensity projection of 500 frames from a video of Cy5-labeled yeast Cdh1 C-terminal IR peptide ((Cy5)- SLIFDAFNQIR) with immobilized APC/C^apo^. **c**, Dwell time distribution from SMOR analysis of a representative movie of the Cdh1 IR peptide binding to APC/C^Apo^ immobilized at the surface.

**Supplementary Data Fig. 6:**
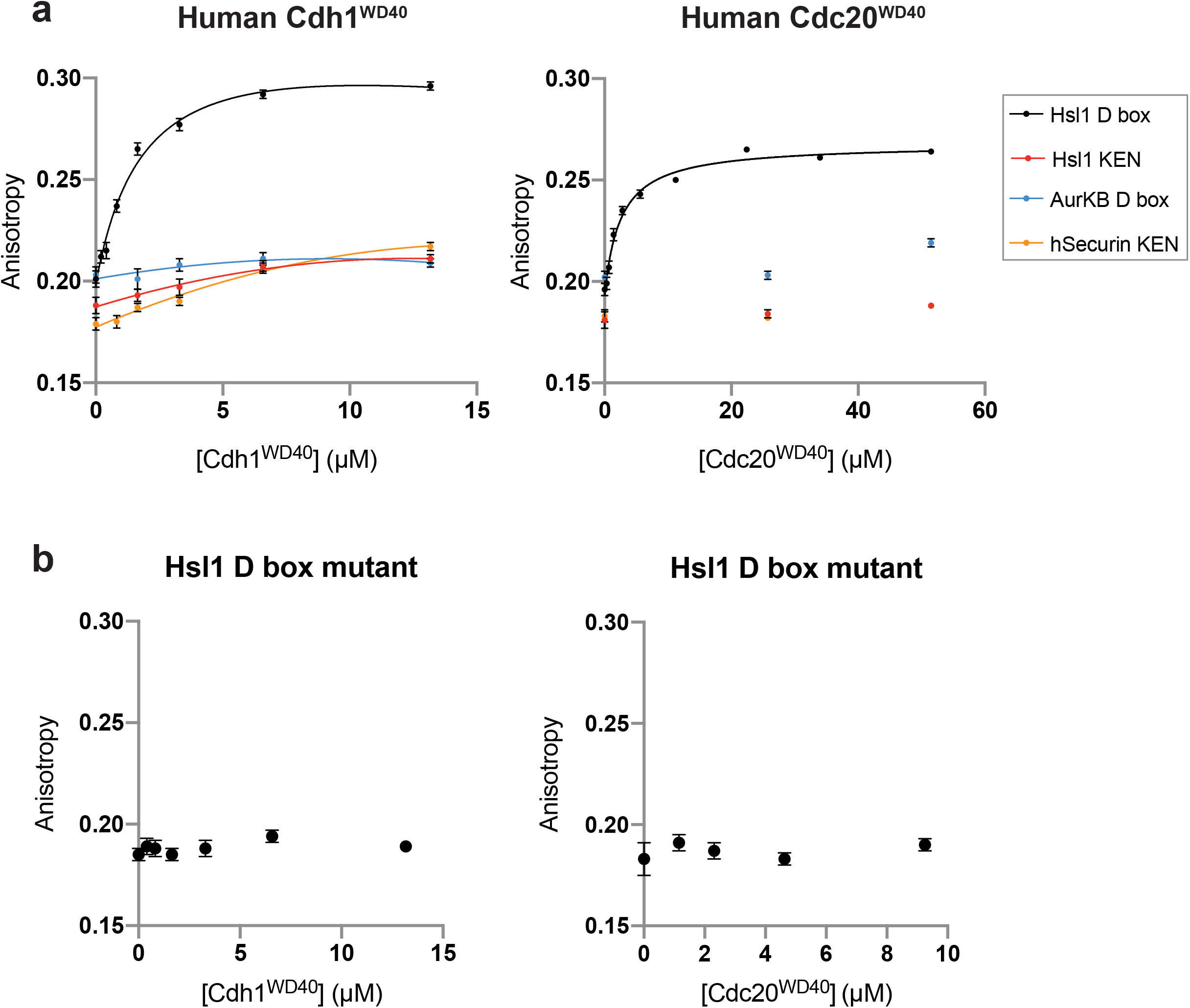
Anisotropy with human activator WD40. **a**, Fluorescence anisotropy analysis of 10 nM Cy-5 labeled peptides (Hsl1 D box, Hsl1 KEN, AurKB D box, and hSecurin KEN) binding to hCdh1^WD40^ or hCdc20^WD40^. **b**, Analysis of mutant Hsl1 D box peptide with hCdh1^WD40^ or hCdc20^WD40^. Data points represent mean +/- SD (n = 10 reads per reaction).

**Supplementary Data Fig. 7:**
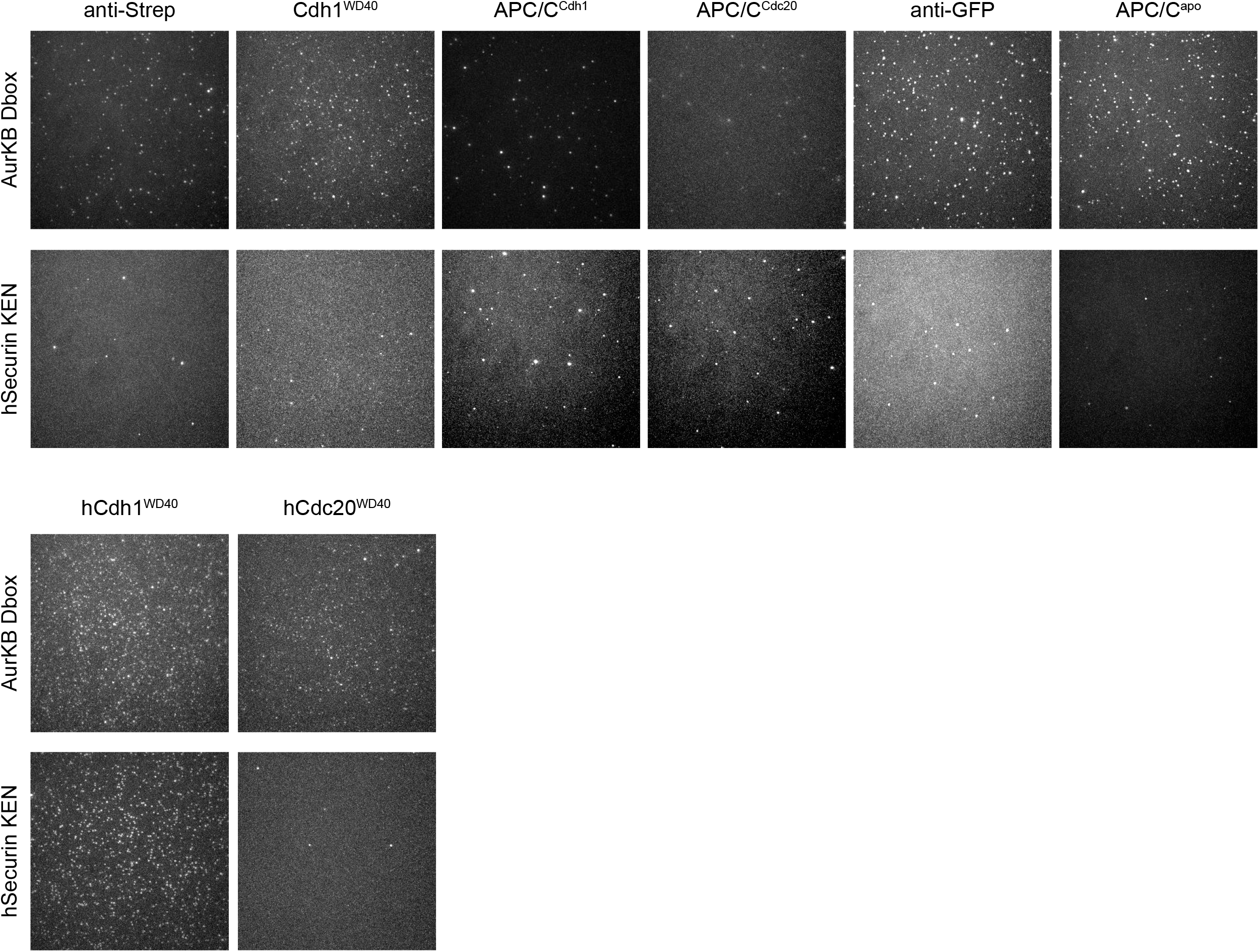
Human degron peptide binding in various conditions. Maximum intensity projections of 500 frames from videos of Cy5-labeled human degron peptides (labeled on left) binding to proteins immobilized at the surface (labeled at top): yeast APC/C and activators in top panels; human activator WD40 domains in bottom panels. Images labeled ‘anti-Strep’ are background controls for binding to APC/C-activator complexes; images labeled ‘anti-GFP’ are background controls for binding to GFP-tagged APC/C^apo^ and GFP-tagged activator WD40 domains.

**Supplementary Data Fig. 8:**
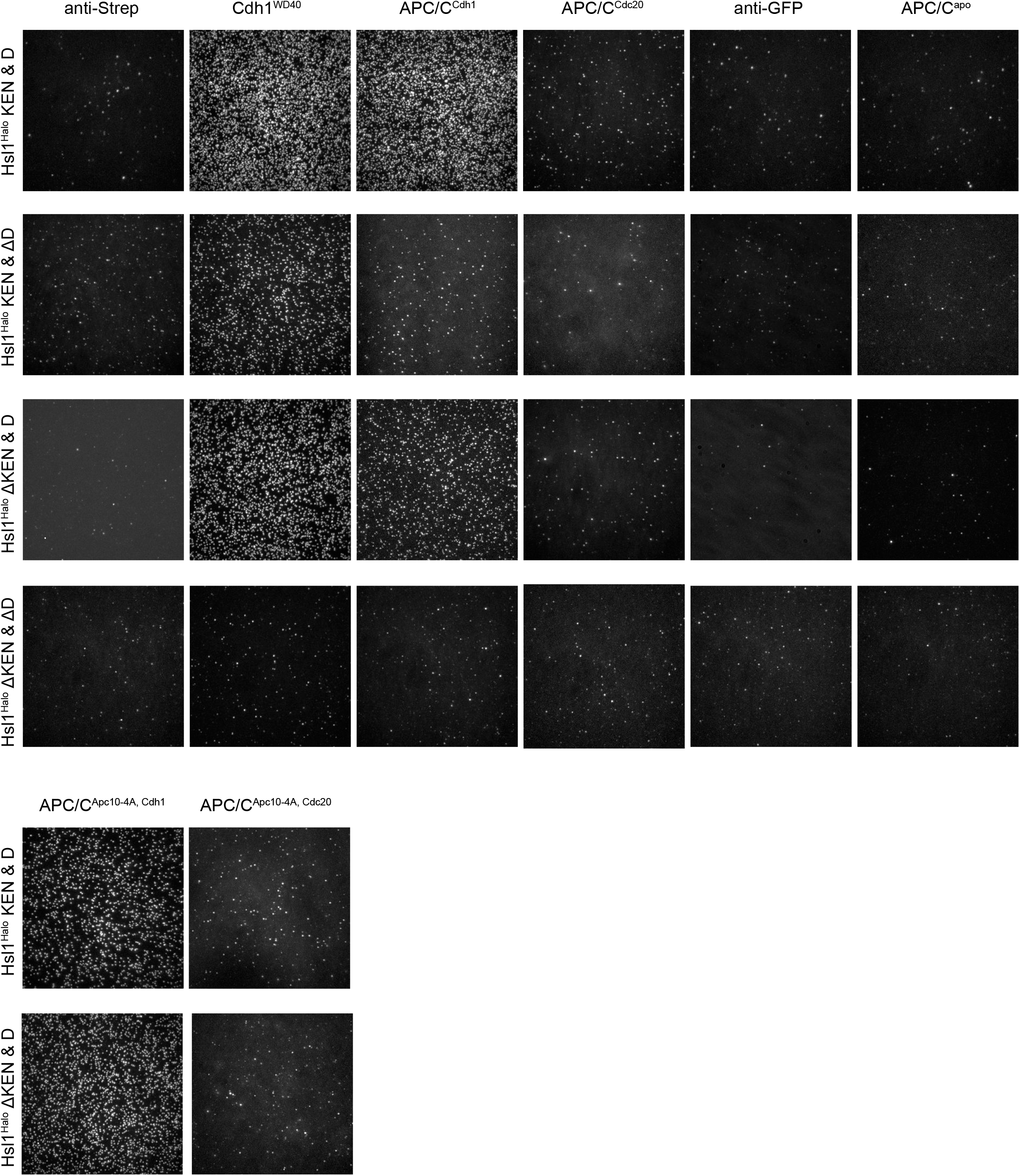
Hsl1^Halo^ binding signal in various conditions. Maximum intensity projections of 500 frames from videos of JF549-labeled wild-type and mutant Hsl1^Halo^ (labeled at left) binding to proteins immobilized at the surface (labeled at top). Images labeled ‘anti-Strep’ are background controls for binding to APC/C-activator complexes; images labeled ‘anti-GFP’ are background controls for binding to GFP-tagged APC/C^apo^ and Cdh1^WD40^.

**Supplementary Table 1:**
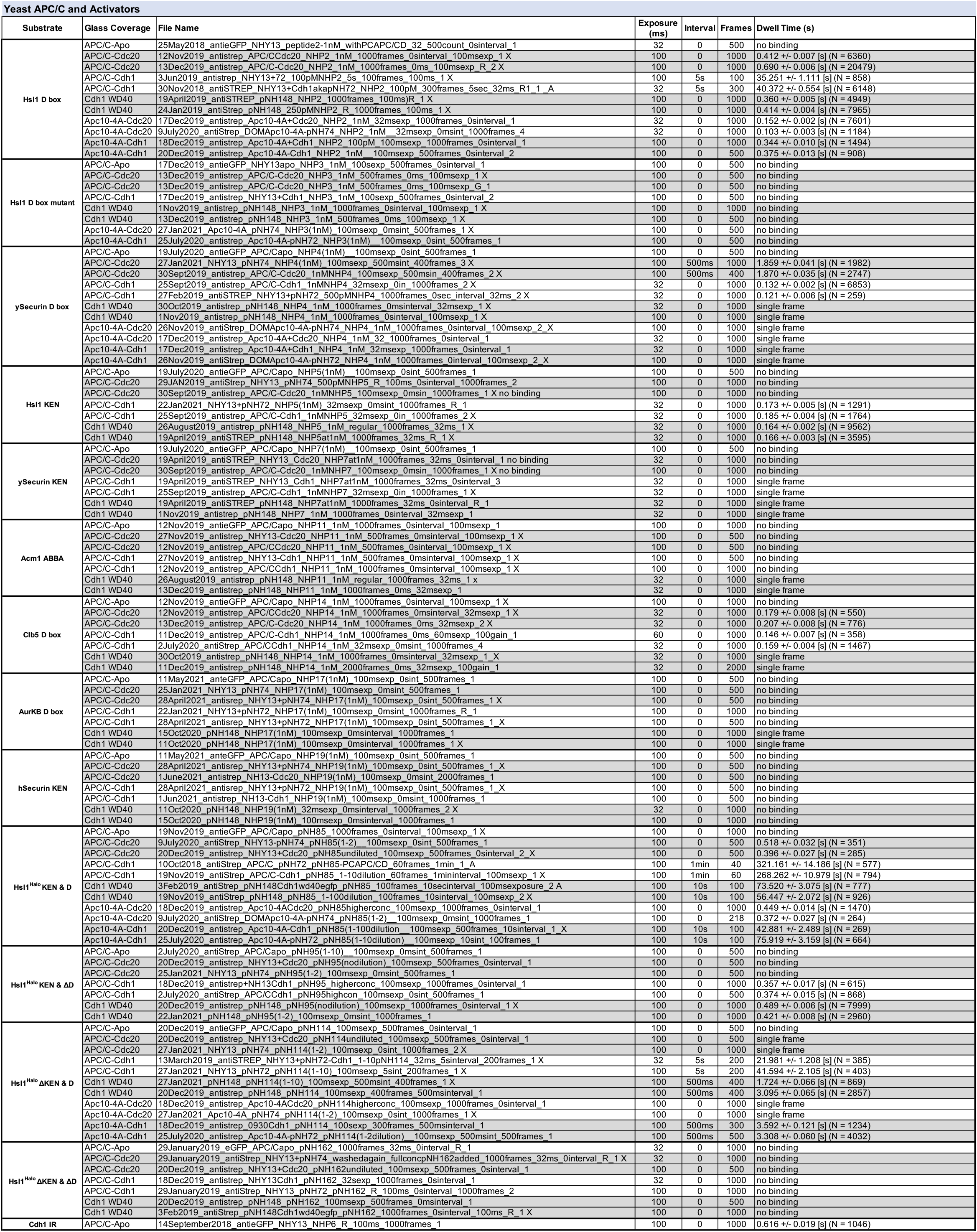
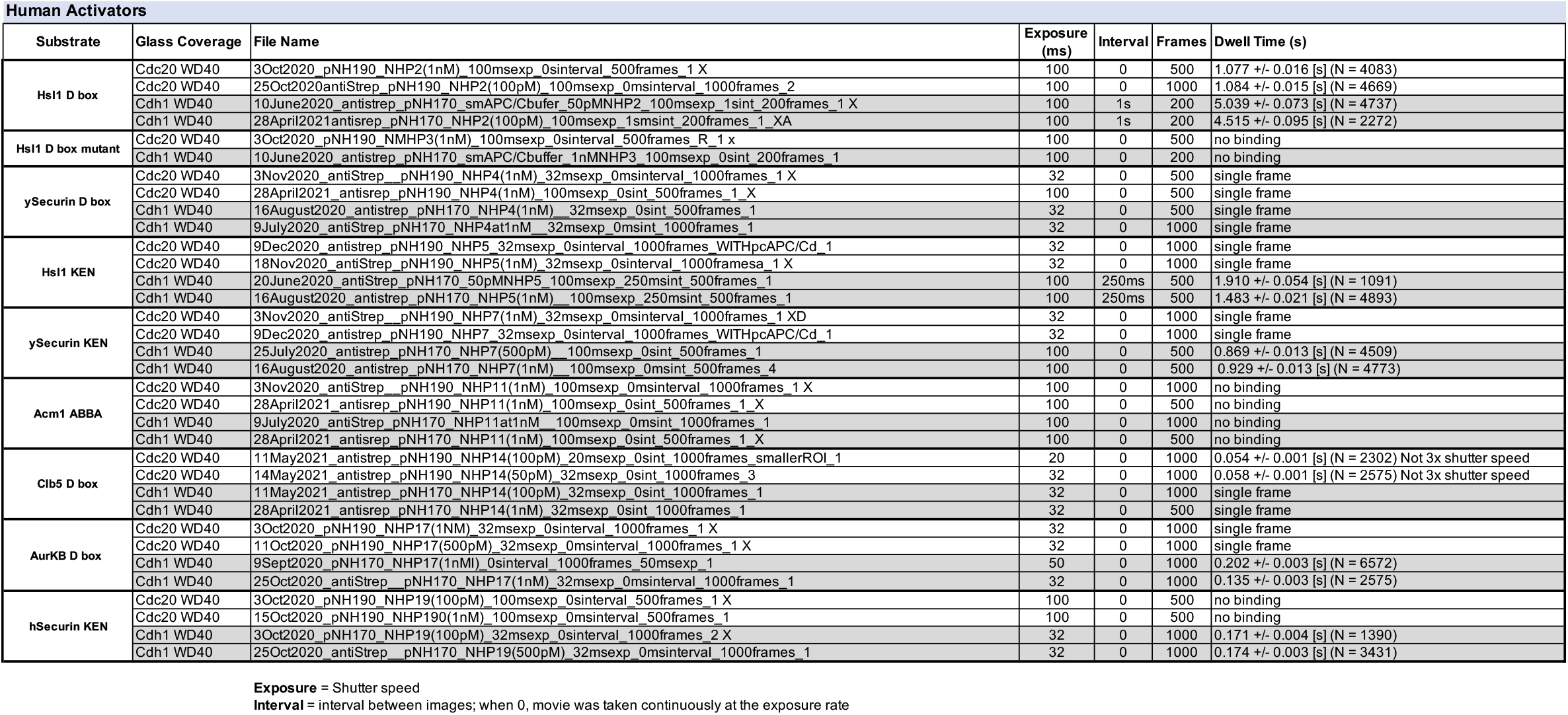
List of representative movies analyzed in this study.

**Supplementary Table 2:** Yeast strains.

|  |  |
| --- | --- |
| NHY13 | cdh1::LEU2<br>bar1::hisG<br>CDC16::CDC16-TAP-HIS3<br>APC1::APC1-GFP-CaUra<br>MATa mating type, W303 background |
| DOM1226 | cdh1:: LEU2<br>bar1::hisG<br>CDC16::CDC16-TAP-HIS3 |
| DOM0930 | CDC16::CDC16-TAP-HIS3<br>doc1d::URA3<br>trp1::TRP1-pRS304-doc1-4A |

**Supplementary Table 3:** Bacmid vectors for protein expression.

|  |  |
| --- | --- |
| pNH72 | 2xStrep-Tag II-yeast Cdh1 |
| pNH74 | 2xStrep-Tag II-yeast Cdc20 |
| pNH144 | 2xStrep-Tag II-yeast Cdh1 WD40 |
| pNH148 | 2xStrep-Tag II-yeast Cdh1 WD40-GFP |
| pNH164 | 2xStrep-Tag II-human Cdh1 WD40 |
| pNH170 | 2xStrep-Tag II-human Cdh1 WD40-GFP |
| pNH175 | 2xStrep-Tag II-yeast Apc10 |
| pNH188 | 2xStrep-Tag II-human Cdc20 WD40 |
| pNH190 | 2xStrep-Tag II-human Cdc20 WD40-GFP |

**Supplementary Table 4:**
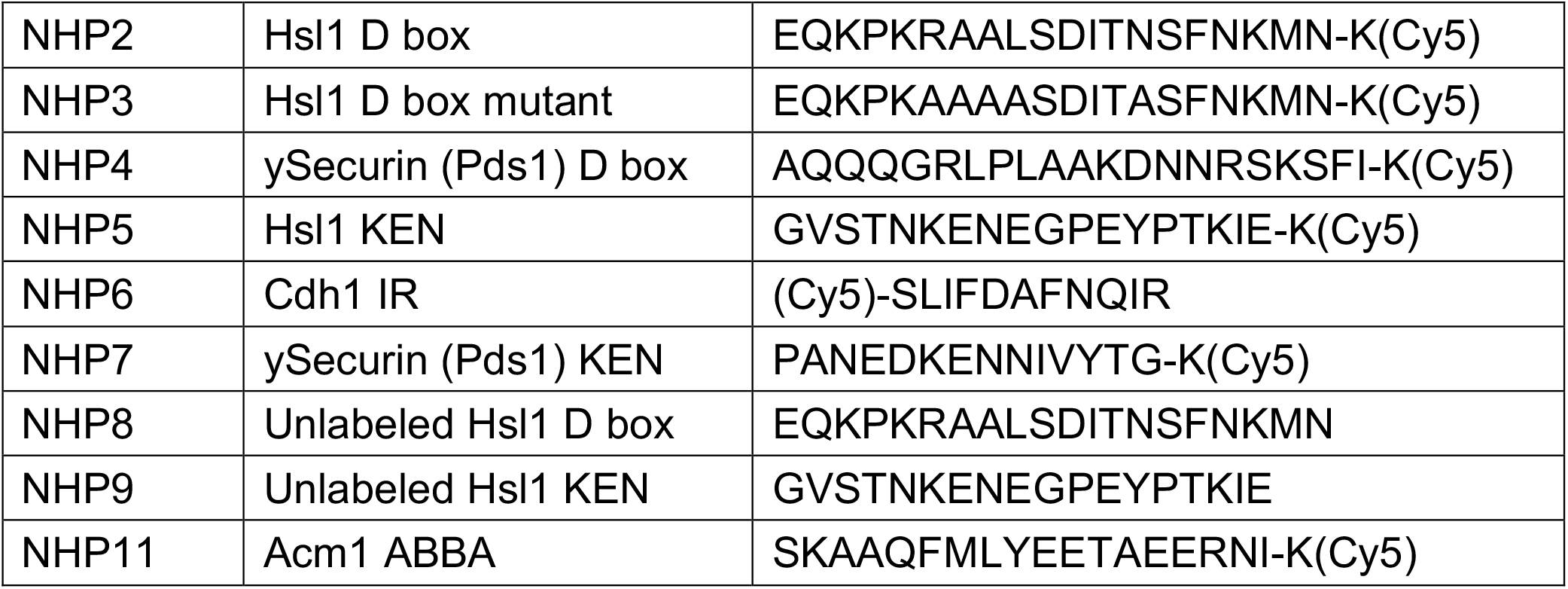
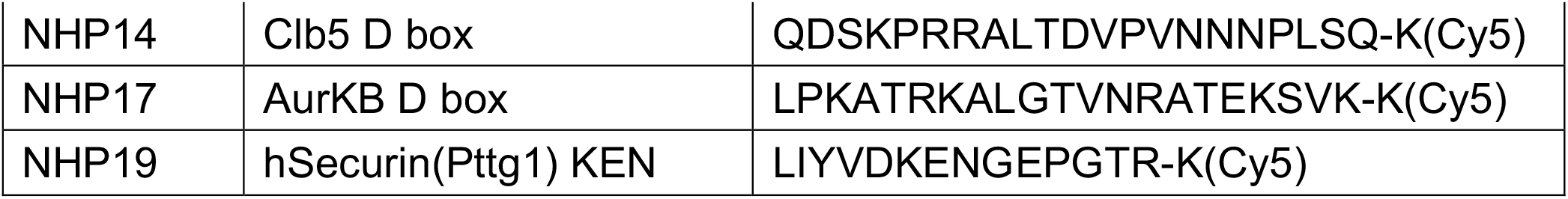
Peptide sequences.

**Supplementary Table 5:** *In vitro* translation plasmids (contain T7 promoter)

|  |  |
| --- | --- |
| pNH85 | Hsl1<sup>667-872</sup>-Halo-TEV-ZZ |
| pNH95 | Hsl1<sup>667-872</sup>-Halo-TEV-ZZ with R,L, & N in D box mutated |
| pNH114 | Hsl1<sup>667-872</sup>-Halo-TEV-ZZ with KEN mutated to AAA |
| pNH162 | Hsl1<sup>667-872</sup>-Halo-TEV-ZZ with KEN & D box mutations |
| pJK567 | hSecurin (Pttg1)-TEV-ZZ |
| pME39 | ySecurin( Pds1)-TEV-ZZ |
| pME60 | Hsl1<sup>667-872</sup>-TEV-ZZ |

## Supplementary Methods: Detailed description of SMOR analysis pipeline

Fluorescent ligand in solution repeatedly binds and unbinds to its binding partner attached to the glass surface. Here, the dwell time during the surface-bound state is determined by the dissociation rate (k_off). When bound, the fluorescent molecule is excited by the TIRF illumination, emits fluorescence that is captured by the camera and appears as a bright spot in the time-lapse movie. In our setup, magnification is adjusted such that each isolated fluorescent spot occupies a ∼3×3 pixel area in the movie. The main goal of the analysis is to extract the dissociation rate from each movie containing many such single molecule binding/unbinding (dwell) events. Here, we assume that all molecules are homogenous and follow the same kinetic rate. In order to achieve this goal, the program identifies the pixel locations where the binding/unbinding events take place, measures the intensity changes over time, identifies the transition between bound and unbound states, aggregates all the dwell events from many molecules at different locations, and calculates the mean dwell time from the dwell time distribution, which is an inverse of the dissociation rate (k_off). As part of the analysis, the algorithm identifies and excludes outliers (such as fluorescent junk or multiple molecules in proximity) and corrects the data (i.e. drift correction and flatfield correction) for better performance. Below are more details of the analysis pipeline, with instructions to allow subsequent users to run the code and adapt the algorithm for their unique experiment/binding data.

Before running the analysis, data needs to be organized in the following manner.

- Data folder

- data1

- movie1.tif
- info.txt
- data2

- movie2.tif
- info.txt

Each data folder must have a single movie file in a tiff file format and an info.txt file that contains the custom parameters for the analysis of each movie. Running a command “run-smor-analysis” starts reading all the movie files under the current working directory and analyzing each movie one by one. Once the analysis is successfully completed, the corresponding figures and a summary file (result.txt) is generated in each data. If an error occurs while analyzing one of the movies, a log file (error.txt) that describes the specific error messages is created in the corresponding data folder.

The analysis pipeline consists of the following steps.

1) Read the movie in a tiff format and the info.txt file.
2) Do flatfield correction.
3) Do drift correction.
4) Find fluorescent intensity peak locations from the maximum intensity Z-projection image.
5) Find the fluorescent spots with similar brightness.
6) For each fluorescent spot trace, apply two-state Hidden Markov Model (HMM) to identify the transition between bound and unbound states.
7) Collect the bound state dwell time from the HMM analysis and calculate the unbinding kinetics.
8) If completed successfully, save the figures and the result file. If not, save an error log file.
9) Move on to the next data folder and repeat.

Description of steps:

### 1) Read the movie and the info.txt file

The code first reads the info.txt file that describes the specifics of the movie and the user-defined custom parameters for analysis. These custom parameters include basic information such as the time interval between frames, spot size in pixels, data filtering conditions, and more. A complete list of all parameters is described at the end. The code uses a python library Tiff file to read the movie in a tiff format. The pixel intensity is loaded in a 3D array (NumPy) where the 1^st^ axis is frame while the 2^nd^ and 3^rd^ are the row and column of each frame image. Then, the image is cropped to 300 x 300 pixels at the center, where the intensity is normally brighter than the edges. If the user finds that the TIRF field is off center, the movie can be cropped to 300 x 300 pixels beforehand.

### 2) Flatfield correction

This code uses the intensity distribution at bright spot locations across the field of view to judge whether to include or exclude those spots. Basically, we want to collect data with similar brightness and exclude outliers. If intensity at a spot is much brighter than other spots, it could be due to multiple spots in proximity or aggregates; hence, they are excluded from the analysis. If intensity is much darker, then it could be fluorescent junk, not valid signal. This approach is valid only if the illumination for the fluorescence signal is uniform across the field of view and the intensity differences is not simply due to non-uniform illumination. In reality, illumination is never completely uniform. In general, the central area is brighter than the outer area due to the Gaussian laser beam profile (Supplementary Data Fig. 4b). Moreover, higher order non-linear patterns might also exist due to aberrations or suboptimal alignment of the optics. Therefore, we used a flatfield correction algorithm to normalize each pixel intensity by the average intensity in the local area. We first make a mask using local binary filter (threshold_local method from skimage library) to identify the pixels where fluorescent spots are located (Supplementary Data Fig. 4b). Then, we obtain the mean intensity within the mask in each bin area (Supplementary Data Fig. 4b). If the bin size is too small, then it reveals a more detailed spatial pattern of illumination, but it becomes less accurate due to a smaller number of mask pixels residing in the bin average. A bin size that is too large yields the opposite effect. We found that a 20 x 20 pixel area works best with our data. If the density of spots is too low, one needs to increase the binning size, and vice versa. The locally averaged image is smoothened a bit using a gaussian filter to remove the sharp contrast between the neighboring bins (Supplementary Data Fig. 4b). The original image is normalized by the smoothened image to compensate for non-uniform illumination (Supplementary Data Fig. 4b). The flatfield-corrected image is once again binned and averaged to double check that the spatial pattern has less structure and looks random (Supplementary Data Fig. 4b). The user can allow or skip this correction by a custom parameter (flatfield_correct = true or false) as needed.

### 3) Drift correction

The microscope sample slide can drift over time and the effect becomes more noticeable if the movie is taken for a long period of time or the microscope setup is less stable. This sample drift can cause problems in our analysis as we use a fixed pixel location to measure the fluorescent intensity changes over time. To solve this problem, the program detects the image drift throughout the frames over time and compensates the amount of drift for every frame. We use an open-source python library for image analysis (*imreg_dft*), using its ‘translation’ method to measure image drift^33^. This method compares two images (test and reference) and finds the translation of the test image in row and column directions as compared to the reference image, using spatial correlation between the two images. In other words, pattern matching with translation is used between the two images. We don’t consider other transformations, such as rotation or magnification, as their effects are less significant than translation. For our analysis, the image at the mid frame was used as a reference as it has the minimum pairwise frame differences with all the other frames; hence, it contains more overlapping spots between the two images. All the fluorescent spots are used as fiducial markers, but only part of them present in both images due to the stochastic nature of binding and unbinding processes. Nevertheless, the algorithm works well if there are sufficient overlapping spots in the two images. Afterwards, additional filters are applied to improve the robustness, since the translation algorithm alone can be affected by unwanted noise, such as bright fluorescent aggregates in some frames. We set a maximum allowed translation between two consecutive frames (default = 2 pixels), followed by a moving average for a lowpass filter, to avoid abrupt jumps between frames, which is more likely due to artifact rather than actual drift since the most dominant drift effect is at low frequency. Drift in both axes can be noted per movie as shown in Supplementary Data Fig. 4c. The user can allow or skip this correction with a custom parameter (drift_correct = true or false) as needed. The need for drift should be determined by the user based on their TIRF microscope. For our microscope, we applied it for acquisition intervals above 500 miliseconds.

### 4) Find fluorescent intensity peak locations from the maximum intensity Z-projection image

After the two previous corrections, we are ready to find the pixel locations where fluorescent ligand binds and unbinds. We use the maximum intensity Z-projection image along the time axis to accumulate all the bright spots from all the frames in the movie. To find the potential binding sites, we use the peak_local_max method from the python library scikit-image, which returns the local maximum peaks from the image. We set a minimum distance between the closest spots allowed to avoid any interference between them. Current default min_distance value is set from the custom parameter (equivalent to spot_size in pixel), so that two peaks cannot be chosen within the area occupied by a single spot. The selected peaks are shown in Supplementary Data Fig. 4d in yellow circles. Subsequently, the average intensity within the spot_size pixel area around each peak is calculated for each frame to measure the intensity changes of the individual spots throughout the frames, and saved as peak_trace. This averaging filter is applied to sum all the signal from each spot around the peak. Some of the peaks are from true binding/unbinding processes, while others could be from inactive fluorescent junk or aggregates of multiple molecules, which will be filtered out in the subsequent steps.

### 5) Find the fluorescent spots with similar brightness

With uniform illumination of fluorescent molecules or with flatfield correction, we can assume that all the fluorescent spots from single isolated fluorophores have similar brightness. Outliers such as very bright signal could be due to multimer and much darker signal could be fluorescent junk. Hence, we use their intensity distribution and cutoff criteria to filter out the outliers. We use both the minimum intensity as well as the maximum intensity distribution as shown in Figure 3c to utilize the information from both bound and unbound states. The maximum (or minimum) intensity value from each peak trace is collected to form a histogram (Fig. 3c). As shown, the intensities are tightly distributed around the peaks. Spots where both their maximum and minimum intensities reside within multiples of standard deviation (SD) from their overall median intensities are selected as inliers (inliers in red and outliers in blue). The multiplication factor is determined by the custom parameters (intensity_min_cutoff and intensity_max_cutoff). For example, cutoff=2 yields the population within two standard deviations from the median intensity. In some experiments, we have observed multiple peaks in the intensity distributions, which indicates that the data includes a mixture of multiple groups. For example, one maximum intensity peak is from normal binding whereas another peak is from transient binding events yielding lower intensity or multimers with brighter intensity. To give the user more options in these scenarios, custom parameters (intensity_min_num, intensity_max_num) are used to set the number of groups or peaks in the intensity distributions, and the parameters (intensity_min_index, ntensity_max_index) are the index of the group we want to include in the analysis. The Gaussian Mixture model from scikit-learn python library was used for the classification of the groups. The selected traces (red) with three parameters (cutoff, num, index) from both min and max intensities are used in the following step to characterize the binding/unbinding kinetics.

### 6) For each intensity trace from the selected spots, apply two-state Hidden Markov Model (HMM) to identify the transition between bound and unbound states

Supplementary Data Fig. 4e is the histogram of intensities along the entire traces of all selected spots. The distribution fits well with the sum of two Gaussian distributions (in red) with means at low and high intensities, which agrees well with our two-states model. Each individual intensity trace shows the transition between unbound state at low intensity and bound state at high intensity, as shown in the example trace figures (Fig. 3d, e). Depending on the experimental parameters, including time interval per frame, total number of frames, and unbinding/binding kinetics, the number of binding/unbinding cycles changes. Fluorescence intensity fluctuates due to several factors, such as intrinsic blinking of the fluorophore, laser power fluctuation, or freely diffusing fluorophores in the background. Hence the true molecular state, either bound or unbound, is hidden under noise and we need to estimate them with a statistical model. A Hidden Markov model (HMM) is a statistical Markov model with hidden states, where the transition probability between states is determined by the current state^34^. We consider a two-state HMM as having only an unbound or single fluorophore-bound state. If the experimental data contains multiple binding states, such as dimers or trimers, the number of states in the HMM needs to be extended (and the code needs to be modified accordingly). For our analysis, we use a python library hmmlearn, and we specifically use the GaussianHMM model as we can assume Gaussian emissions for the intensity fluctuation. HMM uses the entire dataset and tweaks the parameters to maximize the likelihood of observing the experimental data. This means it is robust to the noise sources affecting the state determination, such as blinking or transient spikes, which is superior to other threshold-based methods. For more robust fitting, we use the initial condition of the model parameters from the experimental data, including fractions, means, and variances of the two states from the fitting in Supplementary Data Fig. 4e. Then, the individual trace was fitted into the model to predict the HMM parameters (means, variances, transition probability) as well as the states per frame. After the fitting of the entire traces, we apply additional filters with selection criteria to exclude outliers. Here, we use three criteria based on the mean intensity of unbound states (Supplementary Data Fig. 4f), bound states (Supplementary Data Fig. 4g), and the root-mean-squared-deviation (RMSD) between experimental trace and the trace fitting (Supplementary Data Fig. 4h). We use custom parameters (HMM_RMSD_cutoff, HMM_bound_cutoff, HMM_unbound_cutoff) to select the populations near the peaks within the cutoff multiples of their standard deviations. The traces satisfying all three criteria are finally chosen for the following kinetics analysis. In some experimental conditions, we observed multiple peaks in the bound state intensity distribution, which indicates that there are multiple bound states in the experimental data, such as dark intensity due to transient binding or brighter intensity due to multiple fluorophore binding, regardless of our two states HMM model. In order to give more options to the users in these cases, we use a GaussianMixture model with custom parameters (HMM_bound_num, HMM_bound_index) to select those populations.

### 7) Collect the bound state time from the HMM analysis and calculate the unbinding kinetics

Some of the traces show multiple binding/unbinding events while others have only one or part of the signal (pre-existing or unfinished bound state). To simplify the analysis, we only collect complete bound state times that remain in the bound state from the beginning to the end. First, we collect all the bound state time (i.e. dwell time) from all of the traces. Subsequently, unwanted short or long events can be filtered out. In a short time period, there could be missing events during the sampling or contamination of transient binding events. In a long time period, there could be missing events due to photo bleaching or artifacts due to fluorophores being stuck to the surface. A custom parameter (frame_offset) can exclude short events by offsetting the value from all of the dwell times. Very long dwell outliers that are longer than 10-fold of the median dwell time are automatically excluded from the dwell time collection. Dwell time distribution can be assumed as an exponential distribution with a dissociation rate constant, and in this scenario, the maximum likelihood estimator of the dissociation constant is the inverse of the sample mean dwell time. We can check whether this assumption is valid by comparing the exponential function with the experimental data in the final dwell time histogram where the decay parameter from the experimental sample mean dwell time in red overlays the normalized experimental dwell time histogram in black (as seen in Figure 4a). Our code represents the data as both a probability density function or an inverse cumulative density function. As expected, the data agrees well with the formula in all the dwell time plots reported in this article.

### 8) Save the figures and the result file, if completed successfully, or an error log file

Finally, all of the intermediate steps are saved in figures (png format) and the results are saved in a result.txt file. If an error occurs during any of the steps, a log file error.txt describing the specific error message is automatically created for further investigation or debugging.

#### Info.txt

- time_interval: time interval between frame in seconds
- spot_size: size of diffraction limited spot in pixels
- drift_correct: True or False for drift correction
- flatfield_correct: True for False for flatfield correction
- frame_offset: cutoff in frame number to exclude short dwell events
- save_trace_num: the number of trace figures to be saved as examples
- intensity_min_num: the number of groups in the minimum intensity distribution from the spot traces
- intensity_min_index: the index of group to be selected in the minimum intensity distribution from the spot traces
- intensity_min_cutoff: a multiplication factor to SD to include inliers in the minimum intensity distribution from the spot traces
- intensity_max_num: the number of groups in the maximum intensity distribution from the spot traces
- intensity_max_index: the index of group to be selected in the maximum intensity distribution from the spot traces
- intensity_max_cutoff: multiplication factor to SD to include inliers in the maximum intensity distribution from the spot traces
- HMM_RMSD_cutoff: multiplication factor to SD to include inliers in the RMSD of HMM fitting to the trace
- HMM_unbound_cutoff: multiplication factor to SD to include inliers in the unbound state intensity from the HMM fitting
- HMM_bound_num: the number of groups in the bound state intensity distribution from HMM fitting
- HMM_bound_index: the index of group to be selected in the bound state intensity distribution from HMM fitting
- HMM_bound_cutoff: multiplication factor to SD to include inliers in the bound state intensity from the HMM fitting

#### Code availability

The analysis code is written in Python 3 and available in a Github repository (https://github.com/jmsung/smor-analysis). Follow the instructions in the repository for installation. The code is compatible with several operating systems including Windows, OSX, and Linux. Please contact Jongmin Sung for suggestions or bug reports.

